# Evolution of simple multicellular life cycles in dynamic environments

**DOI:** 10.1101/602250

**Authors:** Yuriy Pichugin, Hye Jin Park, Arne Traulsen

## Abstract

The mode of reproduction is a critical characteristic of any species, as it has a strong effect on its evolution. As any other trait, the reproduction mode is subject to natural selection and may adapt to the environment. When the environment varies over time, different reproduction modes could be optimal at different times. The natural response to a dynamic environment seems to be bet hedging, where multiple reproductive strategies are stochastically executed. Here, we develop a framework for the evolution of simple multicellular life cycles in a dynamic environment. We use a matrix population model of undifferentiated multicellular groups undergoing fragmentation and ask which mode maximizes the population growth rate. Counterintuitively, we find that natural selection in dynamic environments generally tends to promote deterministic, not stochastic, reproduction modes.

## 1 Introduction

The ability of organisms to reproduce is a paramount feature of life, and a great diversity of reproduction modes is observed in nature. Even the simplest organisms, such as colonial bacteria and primitive multicellular species reproduce in various ways: by producing unicellular propagules [Koyama et al., 1977], by fragmentation of the colony into two [Keim et al., 2004] or multiple multicellular pieces [Rippka et al., 1979], and by dissolution of the organism into independent cells [Stein, 1958]. A variety of reproduction modes originates from different external and internal conditions [Bonner, 1998, Strassmann et al., 2000, Rainey and Rainey, 2003, Fiegna and Velicer, 2003, Travisano and Velicer, 2004]. The choice of the reproduction mode has a major impact on the later evolution of the species’ traits. This aspect is especially important for organisms at the brink of multicellular life: the larger the organism grow, the more complex it can become [Smith et al., 2013]. However, it also means longer developmental time, which might incur additional risks; larger propagules require less protection against unfavourable environmental conditions [Shine, 1978], while smaller propagules can be produced in larger quantities [Macarthur and Wilson, 1967, Pianka, 1970]. Thus, the question of the evolution of reproduction modes of simple multicellular organisms and life cycles in general has a paramount importance for our understanding of the history of life on Earth.

Natural selection favours the life cycle, which utilizes the opportunities and handles challenges faced by species the best. The evolution of life cycles among complex organisms is generally slow and with rare exceptions [Sinervo et al., 2000] occurs unnoticed. At the same time, primitive organisms under selection pressure demonstrate an extraordinary ability to adapt their reproductive strategies. Initially, unicellular *Chlamydomonas reinhardtii* experimentally subjected to selection for fast sedimentation in liquid media has evolved into multi-cellular clusters reproducing via single cell bottleneck [Ratcliff et al., 2013a]. Similar experiments with budding yeast *Saccharomyces cerevisiae* show the evolution into snowflake-shaped clusters reproducing by fragmentation [Ratcliff et al., 2013b]. Selection pressure imposed by another source, e.g., the threat from predators, has similar effect [Herron et al., 2019]. It was shown that even prokaryotic unicellular life forms are capable to evolve collective-level traits within a matter of months [Hammerschmidt et al., 2014]. These examples show that natural selection can drive the adaptation of reproductive strategies.

The evolution of life cycles has been investigated from the theoretical perspective as well. Roze and Michod [2001] have studied the evolution of the propagule size and found that the smaller propagules can be selected since they are more efficient in elimination of selfish mutants. Tarnita et al. [2013] considered “staying together” mode of group formation, where cell colonies or organisms grow only by means of division of cells already comprising them (without immigration). There, Tarnita et al. investigated conditions at which the multicellular life cycle characterized by stochastic detachment of unicellular propagules outperforms the unicellular life cycle. Recently, we have performed an extensive investigation for the optimal modes of group reproduction [Pichugin et al., 2017]. It turns out that only life cycles with a regular schedule of reproduction are favoured by natural selection. However, all these studies consider only constant environments, where external conditions do not change with time.

It has been shown that the fluctuations of the environment have a significant influence on the life cycles of many species. Natural examples range from the day-night cycles driving photosynthetic activity [Tamiya et al., 1953], to the spawning of marine invertebrates synchronised with the lunar cycle [Tessmar-Raible et al., 2011, Kaiser et al., 2016], to the alteration of seasons affecting availability of food, energy spendings, amount of daylight, etc [Murphy, 1978, Lenz, 1984, Schierwater and Hauenschild, 1990]. An impact of the environmental changes on the life cycle has also been investigated in evolutionary experiments [Ratcliff et al., 2012, 2013a,b, Hammerschmidt et al., 2014]; and even the changes imposed by human interventions into nature [Gross, 1991] have been reported to have an effect on life cycles.

The hallmark phenomenon observed in dynamic environments is bet-hedging, where organisms combine different reproductive strategies [Philippi and Seger, 1989, Beaumont et al., 2009]. Bet-hedging comes in two flavours: In “between-clutch” bet-hedging, different organisms of the same species use different reproductive strategies. An example of this is the blooming of the succulents which must coincide with a hardly predictable wet season in the desert [Venable, 2007, Gremer and Venable, 2014]. Different plants of the same species have different time of blooming, so those which catch the wet season will successfully reproduce, while others perish. Similar processes occur in many other plants including crops [Silvertown, 1984]. In “within-clutch” bet-hedging, offspring produced together have diverse properties [Einum and Fleming, 2004]. An example is the diversity among egg size in bird clutches: in mild seasons all eggs are hatched, while in harsh seasons only the larger eggs with more nutrients can survive [Olofsson et al., 2009].

A theory describing demographic dynamics in dynamic environments has been developed in [Tuljapurkar, 1989, Orzack and Tuljapurkar, 1989, Tuljapurkar, 1990, Tuljapurkar et al., 2003] for models with discrete time and has focused on random fluctuations of environment. The arising method for the population growth rate is also applicable to continuous time models [Kussel and Leibler, 2005].

As shown in experimental evolution studies [Ratcliff et al., 2013a, Hammerschmidt et al., 2014, Herron et al., 2019], under favourable conditions, the evolution of novel life cycles in microbial populations might occur within a matter of months. Therefore, the ecological and evolutionary processes in fast reproducing populations are intertwined with each other. Especially, environmental fluctuations strongly affect smaller groups because they are more likely to be sensitive to these perturbations in the external environment [Libby and Rainey, 2013, van Gestel and Tarnita, 2017]; small changes in the environment might lead to the significant changes in group behaviour. Yet, the evolution of life cycles of simple multicellularity under dynamic conditions still remains largely unexplored. To which extent can environmental fluctuations affect the patterns of cell colony reproduction? What reproduction modes thrive in dynamic environments? Do dynamic environments enrich the space of life cycles that can evolve, or do they impose additional restrictions? We combine methods from demographic dynamics in dynamic environments with the general framework of fragmentation mode evolution to answer these questions.

## 2 Methods

### 2.1 Life cycle of a group-structured population in a static environment

We consider a population model, where cells are nested into groups. Reproduction of cells leads to growth of groups, but external cells are never integrated into groups (no “coming together” in the sense of Tarnita et al. [2013]). The dynamics of the population is driven by a number of biological reactions representing cell growth and group fragmentation. After each cell division, cells either stay together as a group or fragment [Tarnita et al., 2013, Pichugin et al., 2017]. If they stay together, the group size increases, while the number of groups in the population is unchanged. Such events are given by the reactions

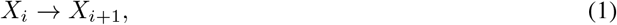

where *X*_*i*_ denotes the group of size *i*. Each cell in a group of i cells has the same birth rate *b*_*i*_, and thus the growth rate of the group is *ib*_*i*_. In our model, birth rates *b*_*i*_ represent the processes influencing the cell growth: they summarise the benefits and costs of a group living within a certain environment. The birth rate of a solitary cell can be set to one (*b*_1_ = 1) without loss of generality. On the other hand, after division, if a group of cells fragments instead of staying together, both the total number of cells and the number of groups in the population increase. The fragmentation results in reactions

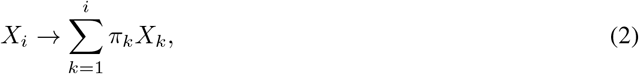

where π_*k*_ is the numer of produced fragments of size *k*. Since fragmentation event conserves the total number of cell, 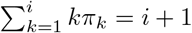.

For example, upon reaching size two after a division from a solitary cell, a group may split into two independent cells, i.e. execute *fragmentation pattern* 1+1, or cells may stay together increasing the group size from 1 to 2. Upon reaching size three, they may fragment into either a bi-cellular group and an independent cell (fragmentation pattern 2+1), three independent cells (fragmentation pattern 1+1+1), or cells may stay together making the group size 3. Upon reaching size four, a group may fragment to one of four fragmentation patterns: 3+1, 2+2, 2+1+1, or 1+1+1+1 (see Fig. 1A), or stay as the group of size 4 and so on. For the sake of calculation efficiency and illustrative purposes, we limit the maximal group size n to 3 in our numerical simulations. However, the approach we developed and our analytical results are not constrained by this limit and are applicable to populations with any group sizes.

**Figure 1:**
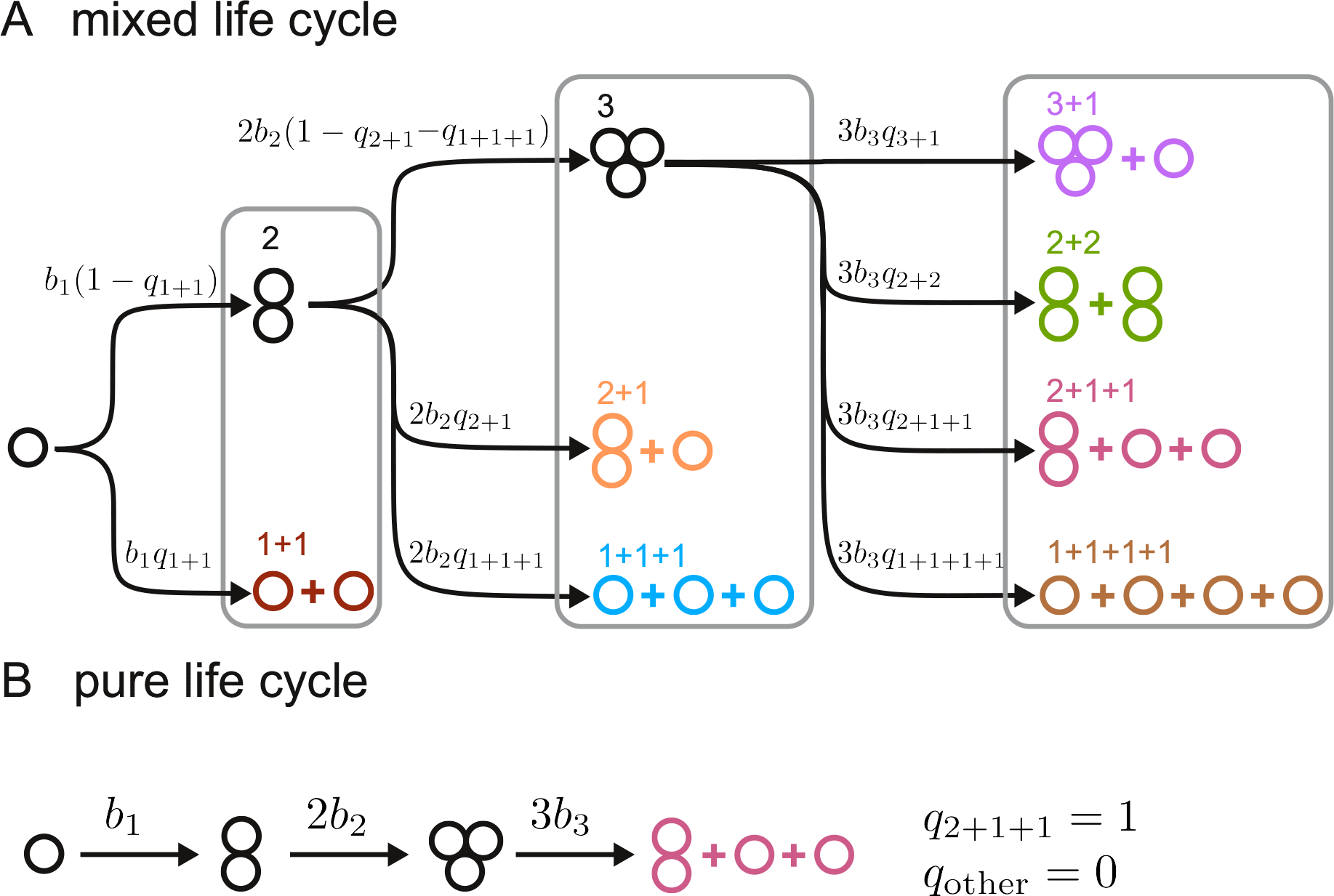
Structure of “staying together” life cycles for a population with the maximal group size *n* = 3. **A**. Schematic figure for the life cycles of a population with *n* = 3. Birth rates *b*_*i*_ are identical for all cells in the same group size *i*. Fragmentation probability set ***q*** = (*q*_1+1_; *q*_2+1_, *q*_1+1+1_; *q*_3+1_, *q*_2+2_, *q*_2+1+1_, *q*_1+1+1+1_) determines the life cycle of the group structure. **B**. Pure life cycles are obtained in a special case where a single fragmentation probability is equal to one, while all others are zero. An example of a pure life cycle using the fragmentation mode **q** = (0; 0, 0; 0, 0, 1, 0) is presented. In a pure life cycle, all groups follow a regular schedule of development and fragmentation.

For an arbitrary life cycle, the rates of reaction are proportional to the probability of the fragmentation pattern κ to occur, denoted as *q*_*κ*_. Thus the set of fragmentation probabilities, ***q*** = (*q*_1+1_; *q*_2+1_, *q*_1+1+1_; *q*_3+1_, *q*_2+2_, *q*_2+1+1_, *q*_1+1+1+1_; …), defines the *fragmentation mode* of a population. In the special case where only a single fragmentation reaction occurs, i.e. all fragmentation probabilities except one are zero, the life cycle represents a regular schedule of the group development and fragmentation. We refer to such cases as *pure life cycles*, see Fig. 1B. In other cases, commonly referred to as *mixed life cycles*, the sum of fragmentation probabilities at each size cannot exceed one: *q*_1+1_ ≤ 1 and *q*_2+1_ +*q*_1+1+1_ ≤ 1; at the maximal size it has to be one, so in our simulations we used *q*_3+1_ + *q*_2+2_ + *q*_2+1+1_ + *q*_1+1+1+1_ = 1 (see Fig. 1A).

The rates of reactions are density independent and therefore, lead to a set of linear differential equations describing the population dynamics, see Appendix A and [Pichugin et al., 2017] for details,

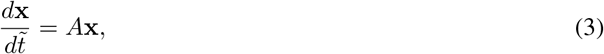

where **x** is the vector for abundances *x*_*i*_ of groups size *i*, 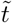 is time, and *A* is the projection matrix. In a static environment, the projection matrix *A* does not change over time. Therefore, the population dynamics converges to a stationary regime where

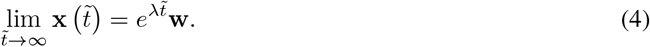

Here, λ is the leading eigenvalue of the projection matrix *A*, and **w** is the right eigenvector associated with λ. The growth rate of the population is determined by λ expressed in terms of birth rates *b*_*i*_ and the fragmentation mode **q**. Hence, evolution favours the life cycle which gives the largest λ. For a static environment, it has been shown that an evolutionarily optimal life cycle must be a pure binary life cycle, where a parental group fragments into exactly two offspring groups [Pichugin et al., 2017].

### 2.2 Growth of the group-structured population in a dynamic environment

Next, we investigate the evolution of life cycles under dynamic environmental conditions, where growth rates *b*_*i*_ do not remain the same through time. Here, we consider a dynamic environment in a form of a regular switch between two seasons 𝑆_1_ and 𝑆_2_. Each season is characterized by its own set of birth rates: 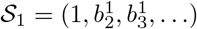 and 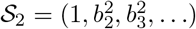, respectively. Consequently, 𝑆_1_ and 𝑆_2_ may favour different life cycles. In this setting, 𝑆_1_ lasts for time duration τ_1_ and then switches to 𝑆_2_, which lasts for τ_2_. Hence, the dynamic environment is determined by the two sets of birth rates and two season lengths *𝒟* = {𝑆_1_, τ_1_; 𝑆_2_, τ_2_}.

In the dynamic environment, the growth rate of the population cannot be characterized by a single projection matrix. However, the demographical dynamics within a single season is still described by a single projection matrix. Therefore, we numerically simulated the population growth in a dynamic environment using the corresponding projection matrix during each season. We follow the method used in [Kussel and Leibler, 2005] to compute the average population growth rate (Λ) in a dynamic environment (*𝒟*) over a whole sequence of seasons, as a slope in the logarithm of the populations size against time, see Fig. 2 and Appendix B. The average growth rate Λ in a dynamic environment plays the same role as the leading eigenvalue λ of the projection matrix in a static environment: the life cycle with higher Λ will eventually outgrow others with lower Λ.

**Figure 2:**
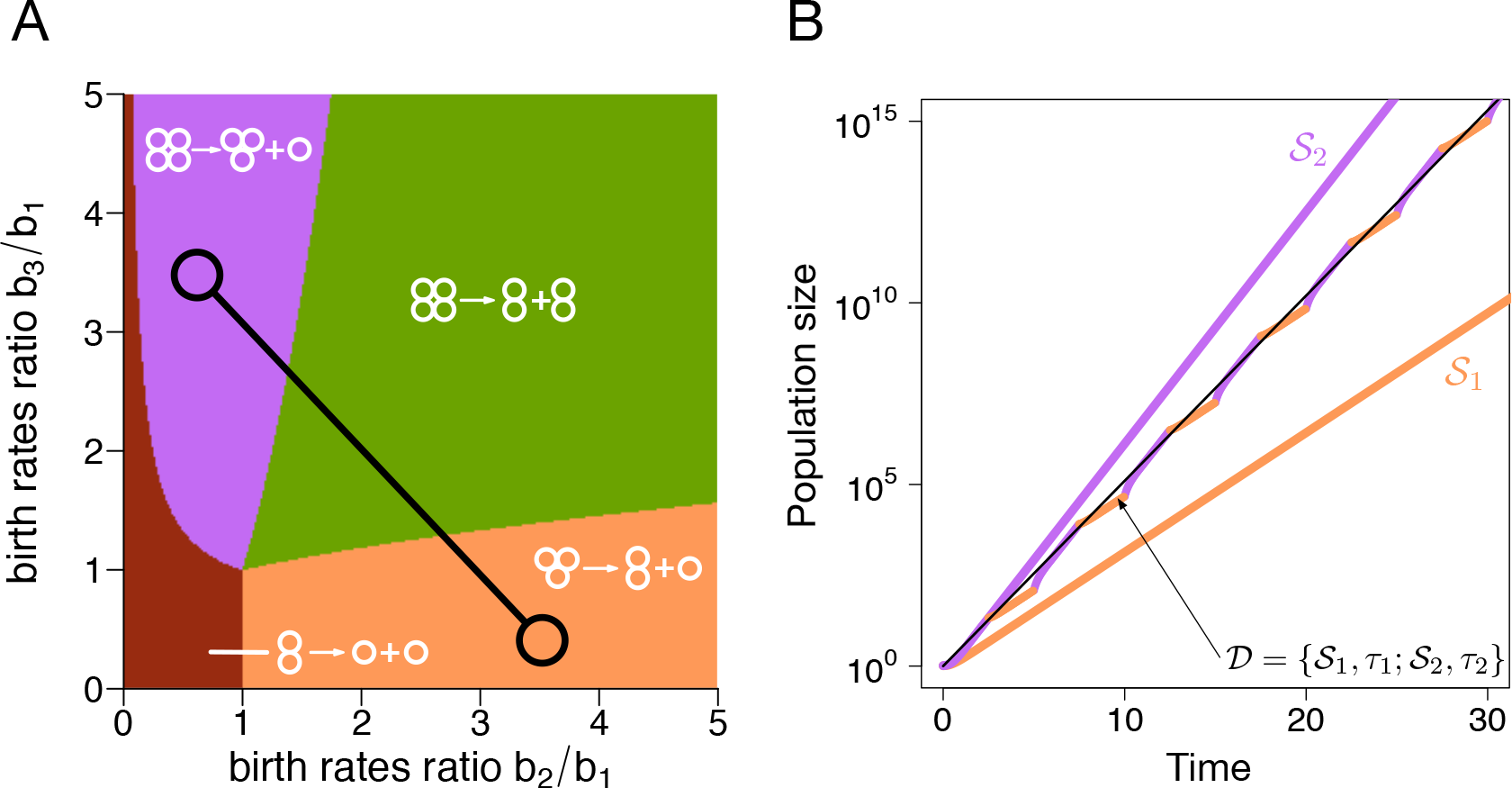
Unconstrained population growth in dynamic environment can be approximated by exponential growth. **A**. The map of evolutionarily optimal life cycles in static environments. Only four pure binary life cycles are evolutionarily optimal: 1+1, 2+1, 3+1, or 2+2. The dynamic environment with seasons 𝑆_1_ and 𝑆_2_ is represented by a pair of interconnected circles. **B**. The growth of population executing a pure life cycle 3+1 (**q**_3+1_ = (0; 0, 0; 1, 0, 0, 0)) in dynamic and static environments. Each line shows the temporal growth of the population size. Coloured lines correspond to the population growth in static environments 𝑆_1_ = (1, 3, 0.5) and 𝑆_2_ = (1, 0.5, 3), respectively. The two-colored line indicates the population growth in the dynamic environment, alternating between two seasons 𝑆_1_ and 𝑆_2_ with *τ*_1_ = *τ*_2_ = 2.5. While the growth in the dynamic environment is complicated in general, it can be approximated very well by exponential growth (thin black line).

For each studied dynamic environment *𝒟*, we numerically find evolutionarily optimal life cycles by maximizing Λ(**q**, *𝒟*) with respect to the vector of fragmentation probabilities **q**, see Appendix C. Note that we perform optimization on a multi-dimensional lattice of **q**, so an accuracy of the optimal life cycle **q** is limited by the lattice spacing, which we set to 0.05. We repeat the optimization for different initial conditions to take into account the possibility of multiple local optima.

## 3 Results

### 3.1 Limit regimes of dynamic environments

For a given pair of seasons {𝑆_1_, 𝑆_2_}, we screened a wide range of seasons lengths combinations {τ_1_, τ_2_} and found a set of locally optimal life cycles for each dynamic environment *𝒟* = {𝑆_1_, τ_1_; 𝑆_2_, τ_2_}. We present the results of this screening in the form of optimality maps, which indicate optimal life cycles at a given set of season lengths. Each pixel on a map represents a set of season lengths {τ_1_, τ_2_}, and the colour of a pixel is given by the all optimal life cycles found in many optimizations from different initial conditions, see Appendix D. For convenience, we convert parameters τ_1_ and τ_2_ into the season turnover period *T* ≡ τ_1_ + τ_2_ and the ratio of season lengths *t* ≡ τ_1_/τ_2_ and present the obtained map using *T* and *t*. Examples of optimality maps are presented in Fig. 3.

**Figure 3:**
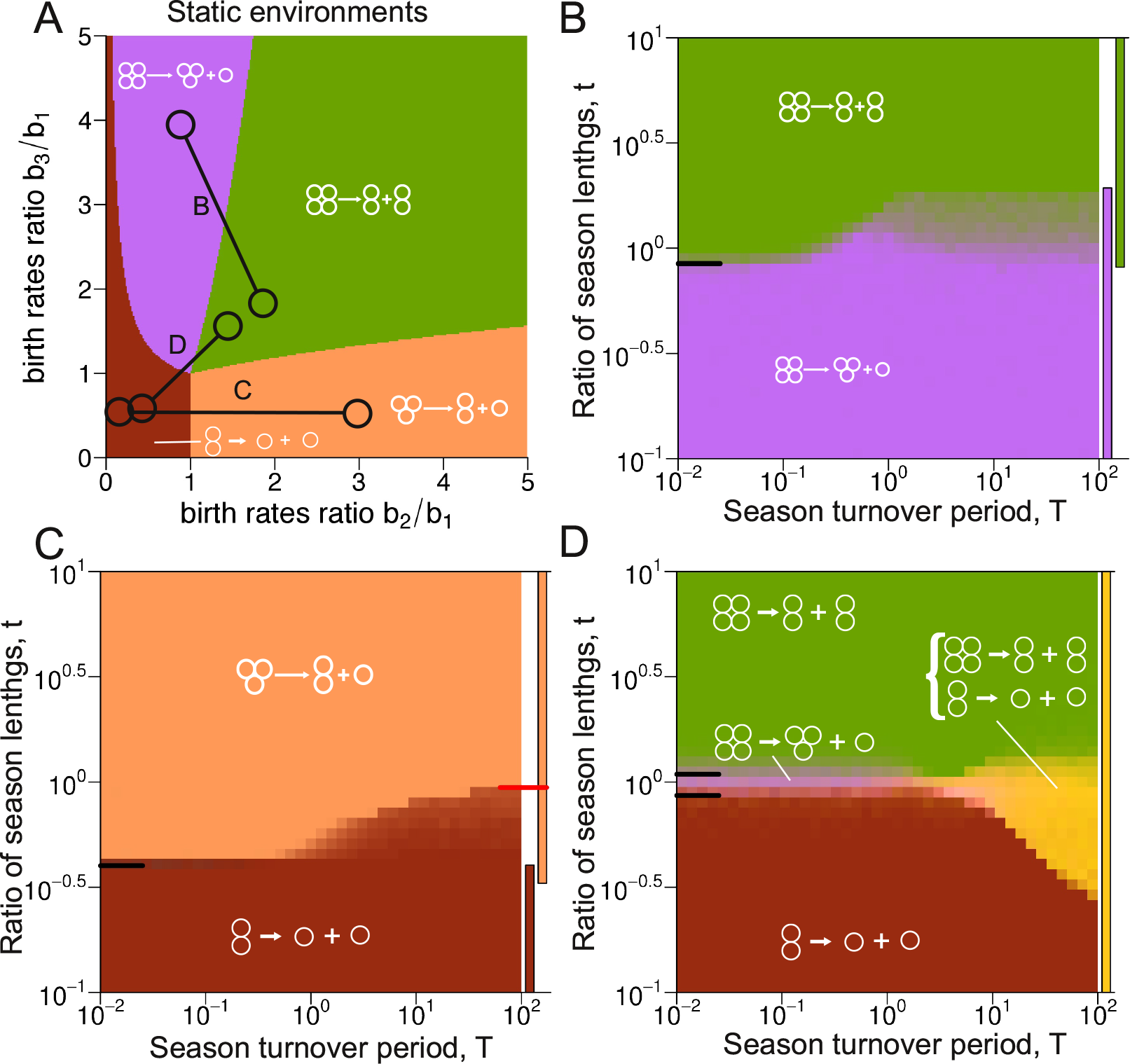
Evolutionary optimality of life cycles in extreme regimes can be described by analytical approximations. **A** The dynamic environment used in panels B, C, and D are represented by connected circles. **B** The optimality map featuring only pure life cycles. When season 1 is much longer than season 2, *t* ≫ 1, regardless of the overall seasons turnover length, the optimal life cycle is always the pure life cycle 2+2. For the opposite case, *t* ≪ 1, 3+1 is always optimal. For short seasons, *T* ≪ 1, the border between two life cycles is located at the position predicted by the short seasons approximation (black tick line on the left side of the map). For long seasons *T* ≫ 1, for this dynamic environment, the transition between two life cycles is performed through the bi-stability between pure life cycles, as suggested by long seasons approximation (see main text). Coloured bars on the right side of the map show the areas of stability of pure life cycles according to the long seasons approximation. **C** The optimality map featuring a mixed life cycle at the long seasons regime. For *t* > 0.287, pure life cycle 1+1 is not evolutionarily optimal, yet the mixed life cycle featuring pattern 1+1 persists up to *t* = 0.485, where it disappears in a saddle-node bifurcation indicated by red mark (see main text). **D** The pure life cycles 1+1 and 2+2 can be executed within the same mixed life cycle at different seasons (see main text), and such a combination (yellow) might be evolutionarily optimal in long seasons regime. Exact parameters used in calculations and simulations are presented in Appendix L.

We focus our analysis on extreme regimes: prevalence of the the single season (*t* ≪ 1 or *t* ≫ 1), short seasons (*T* ≪ 1), and long seasons (*T* ≫ 1). The numerical simulations show that at intermediate lengths of seasons (*t* ∼ 1 and *T* ∼ 1), the behaviour of the system is intermediate between these extremes, see Fig. 3.

The simplest behaviour occurs in the prevalence regime (*t* ≪ 1 or *t* ≫ 1). In this case, the influence of the shorter season on the population growth is negligible. The growth rate in such dynamic environment is close to the growth rate in the static environment given by the long season, i.e. Λ(**q**, *𝒟*) ≈ λ(**q**, 𝑆_1_) when the first season is much longer (*t* ≫ 1). As a consequence, a pure binary life cycle, which is optimal in the static environment provided by the first season 𝑆_1_, is evolutionarily optimal in the dynamic environment *𝒟*, where the first season prevails (*t* ≫ 1). Similarly, the life cycle optimal in the static environment provided by the second season 𝑆_2_ is evolutionarily optimal in dynamic environments, where *t* ≪ 1. All numerically obtained optimality maps confirm this.

For short seasons (*T* ≪ 1), the population composition changes little within a single period of seasons change. Thus, demographical changes in the population occur at much slower time scale than changes of the environment. As such, the system effectively experiences the average environment with season lengths being the weights of each component [Gokhale and Hauert, 2016]. Thus, in the *short seasons approximation* — the population growth rate is given by the growth rate in the averaged static environment 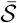

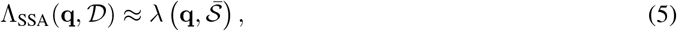

where

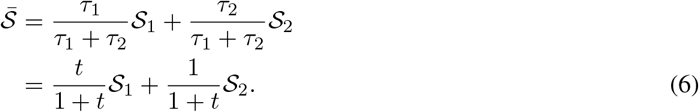

This approximation also allows us to use the results of the optimal life cycles in a static environment [Pichugin et al., 2017]. These imply that for any dynamic environment with short seasons, there is only a single evolutionarily optimal life cycle, which is a pure binary life cycle. In addition, the short seasons approximation allows us to explicitly find the border value of *t* separating the area of optimality of favoured life cycles in the static environment given by 𝑆_1_ and 𝑆_2_, respectively. The border is determined by the ratios of season lengths *t* at which the average environment lies at the border between areas of optimality in static environments. Numerical simulations reproduce this border well, see the predicted border marked at the left sides of the optimality maps in Fig. 3.

For the long season length (*T* ≫ 1), the population reaches the stationary regime within each season, and the transient growth regime between two adjacent seasons can be negligible. This suggest the *long seasons approximation* — the population growth rate is given by the weighted average of growth rates in static environments

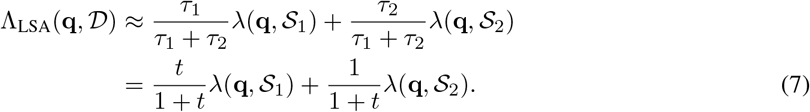

Under the long seasons approximation at intermediate values of t, the optimal life cycle is not necessarily pure, and there might be more than one locally optimal life cycle, see examples on Fig. 3C. The transition in *t* from the prevalence of one season to another is non trivial in this case. These changes in optimal life cycles along *t* in the long seasons regime (*T* ≫ 1) are our focus in the remaining part of this study.

### 3.2 Borders of the prevalence regimes

In the limits *t* → ∞ and *t* → 0, the optimal life cycles are determined by the prevalent season. However, as we increase the other season length, the optimal life cycle may change. In this section, we examine the border of these prevalence regimes. We begin by considering extremely small *t* and measure how long the optimal life cycle persists against the increase of *t*.

We sampled 40, 000 pairs of seasons and investigated at which *t* the prevalence regime is violated, see Appendix E for details of sampling. We found that the border of the prevalence regime significantly varies between season combinations. For some pairs of seasons, the prevalence regime is extremely robust while some others show that the prevalence regime is extremely fragile; even at *t* = 10^−4^ the prevalence was violated.

We found that the main factor determining the robustness of the prevalence regime is the environment in the prevalent season. The most fragile prevalence regimes were observed for environments at the border between two different types of life cycles and environments promoting unicellular life cycle, see Fig. 4A. At the same time, the most robust prevalence regimes were observed for environments far from optimality borders, see Fig. 4B.

**Figure 4:**
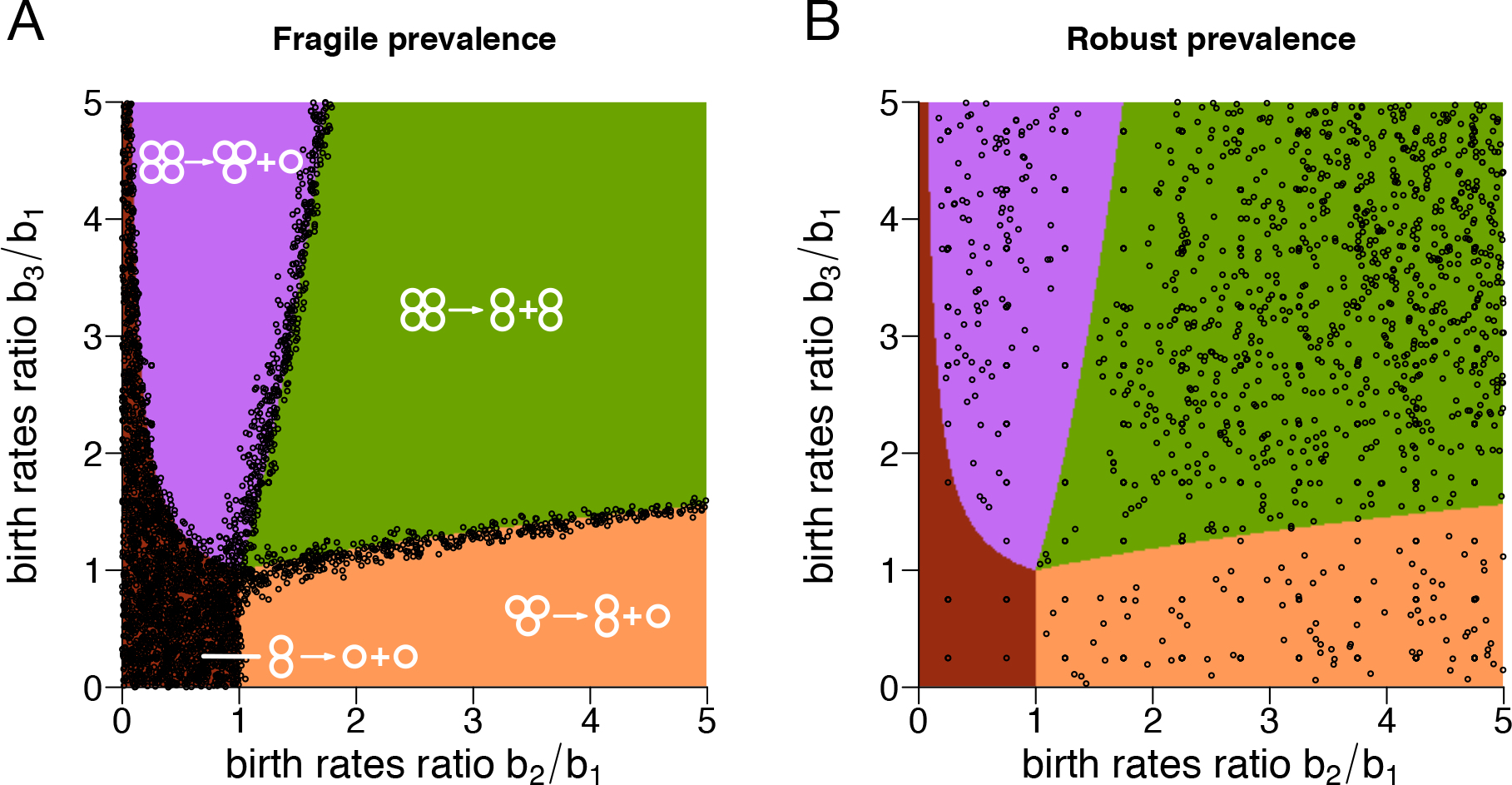
The robustness of the prevalence regime is mainly determined by the position of the prevalent environment. **A**. Circles show prevalent seasons, for which the violation of the prevalence regime occurred already when the fraction of time *t* spend in the second, randomly chosen environment is small, *t* < 0.1. These non-robust seasons either promote unicellularity or are located at the borders between areas of optimality - where at least two life cycles have similar growth rate. **B**. Circles show prevalent seasons, for which the violation of the prevalence regime occurred only when the fraction of time spend in the other environment, is very large, *t* > 10. The distribution of these seasons has low density in the border regions.

### 3.3 Stability of life cycles in the long seasons regime

In this section, we investigate what kind of life cycles emerge to be evolutionary optimal at intermediate *t* in the long seasons regime (*T* ≫ 1). In the long seasons regime, the population growth in a dynamic environment can be inferred from the growth rates in two stationary environments, see Eq. (7). Therefore, the analysis of evolutionary optimality of life cycles can be performed with relatively simple expressions.

An arbitrary fragmentation mode (**q**) is a local optimum of the growth rate Λ, when any small change in the probabilities set **q** leads to a decrease in the population growth rate. For this to happen, all fragmentation patterns κ must fulfil the conditions

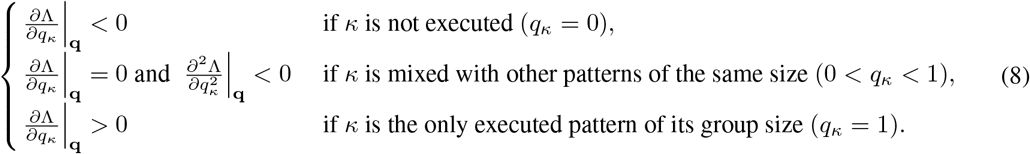

First, we consider the evolutionary optimality of pure life cycles, where only one fragmentation pattern occurs, see Fig. 1B. These life cycles establish a regular schedule of group growth and reproduction, which is commonly observed in nature. From the perspective of our model, in pure life cycles *q*_*κ*_ = 0 or *q*_*κ*_ = 1, the investigation of evolutionary optimality invokes only the first order derivatives, see Eq. (8). A pure life cycle **q** becomes evolutionary unstable when an admixture of at least one of absent fragmentation patterns (with *q*_*κ*_ = 0) no longer decreases the growth rate: 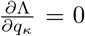. For a given pair of the pure life cycle **q**_1_ and admixture life cycle **q**_2_, this is achieved at the ratio of the season length

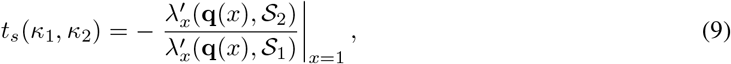

where **q**(*x*) is the mixed life cycle in which fragmentation occurs by *κ*_1_ with probability *x* and occurs by *κ*_2_ with probability 1 − *x*, see Appendix F for details. Knowing the values of *t*_*s*_ for all pairs (*κ*_1_, *κ*_2_) makes it possible to outline the ranges of *t* where each pure life cycle is optimal. We denote this optimality areas as coloured bars to the right of each optimality map, see Fig 3.

Next, we consider mixed life cycles, which can emerge as evolutionary optimal in the long seasons regime. Above, we have shown that only pure life cycles are evolutionary optimal if there is effectively a single season (*t* ≪ 1 or *t* ≫ 1). Therefore, as *t* approaching these extreme values, evolutionary optimal mixed life cycles cease to exist. This happens by one of two scenarios: either a mixed life cycle transforms into a pure one, or it merges with the local minimum of Λ and disappears in a saddle-node bifurcation. The majority of mixed life cycles observed in our simulations feature only two fragmentation patterns. For these life cycles, transition from a mixed optimal life cycle into a pure one occurs at values *t* given by Eq. (9). The saddle-node bifurcation (if exists) occurs at *x* = *x*^∗^ and *t* = *t*^∗^ satisfying

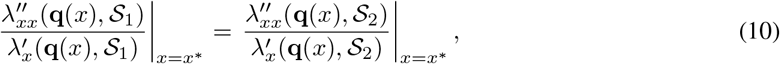

and

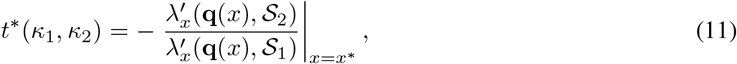

see Appendix G for detailed analysis of mixed life cycles optimality. We highlight the positions of saddle-node bifurcations *t*^∗^ by red marks on the right hand side of our optimality maps, see Fig 3C.

Finally, we found a distinct solution, when the maximal size of groups produced in one fragmentation pattern is smaller than the minimal size of offspring produced by another pattern. With the maximal group size 3, there is a single such pair: *κ*_1_ = 1+1 and *κ*_2_ = 2+2. When the two seasons in the dynamic environment favour the pure fragmentation modes 1+1 and 2+2, the optimal life cycle in a dynamic environment in a long seasons regime is always **q**_*C*_ = **q**_1+1_ + **q**_2+2_ = (1; 0, 0; 0, 1, 0, 0), see Appendix I for details. In the long seasons regime, a population employing **q**_*C*_ is capable to execute pure life cycle 1+1 during seasons favouring 1+1 over 2+2; and pure life cycle 2+2 during seasons favouring 2+2 over 1+1. We call such scenario a coexisting fragmentation mode **q**_*C*_ and the optimality map in Fig. 3D presents this. Except in this special case, numerical simulations confirm our analytical results.

### 3.4 The spectrum of evolutionary optimal life cycles in dynamic environments is diverse

In this section, we consider, which fragmentation patterns can contribute to evolutionarily optimal life cycles. We begin our analysis from a specific scenario, where all birth rates are similar to each other. From a technical point of view, the situation where all birth rates are equal, constitutes a neutral environment at which all growth rates of any mixed or pure life cycles are equal. We consider near neutral environments, where cell birth rates in both seasons slightly deviate from one *b*_*i*_ = 1 + εβ_*i*_, ε ≪ 1, where β_*i*_ and thus *b*_*i*_ is different in the two seasons. As a consequence, in the vicinity of this neutrality point (ε ≪ 1), the growth rate of any life cycle is close to one and can be represented in a form Λ(**q**, *𝒟*) = 1 + εΛ_1_ + *O*(ε^2^), where Λ_1_ is associated with the first derivatives, see Appendix J. Biologically, this corresponds to a scenario, where living in a group has only minimal impact on the cell growth. This scenario is seems to be relevant in the early stages of the evolution of multicellularity, where benefits of the group formation are minimal due to the absence of adaptations to collective life.

There, the case where the maximal group size is limited to two is analytically trackable, see Appendix J. For groups not exceeding size two, only three fragmentation patterns are available: 1+1 (unicellularity), 2+1 (binary fragmentation) and 1+1+1 (multiple fragmentation). We proved that only three types of life cycles can emerge in near neutral environments: pure unicellularity and binary fragmentation, as well as the mixed life cycle utilizing both of them, see Fig. 5. The multiple fragmentation pattern is unable to contribute to evolutionarily optimal life cycles here.

**Figure 5:**
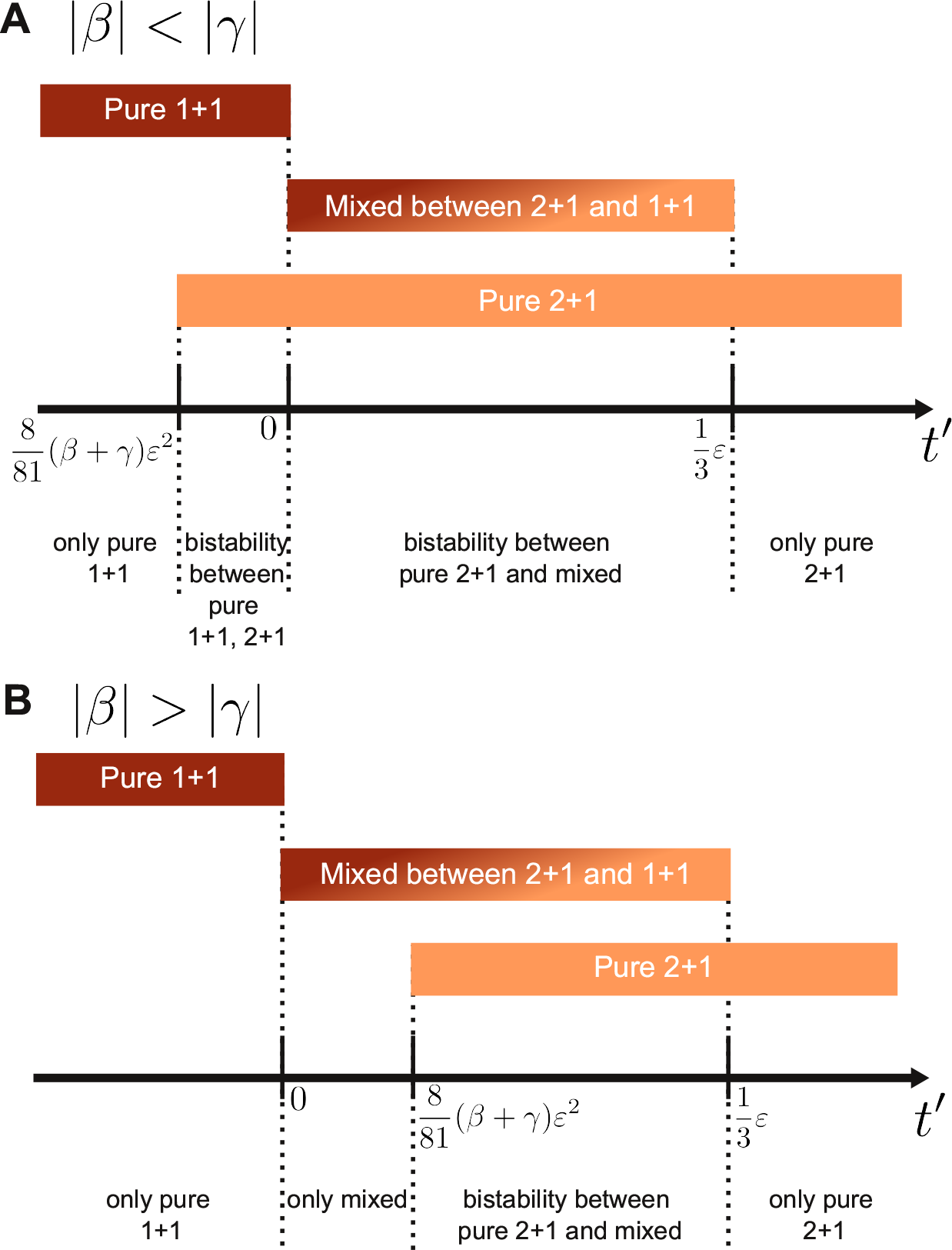
Only three kinds of life cycles can be evolutionarily optimal in near neutral environments if the maximal group size is *n* = 2. Populations, in which group size do not exceed two, have an access to three fragmentation patterns: 1+1, 2+1 and 1+1+1. In near neutral environment constructed by 𝑆_1_ = {1, 1 + *εβ*} and 𝑆_2_ = {1, 1 + *εγ*}, only three fragmentation modes can be evolutionarily optimal: pure 1+1, pure 2+1, and a mixed life cycle simultaneously utilizing fragmentation modes 1+1 and 2+1. The fragmentation mode 1+1+1 does not contribute to an evolutionary optimum under any near neutral dynamic environment. Since the same signs for both *β* and *γ* give the same optimal life cycle in both seasons, we focus on different signs: *β* > 0 and *γ* < 0. **A** For |*β*| < |*γ*|, there is the range of *t* where population exhibits a bi-stability between two pure life cycles. **B** For |*β*| > |*γ*|, there is the range of *t* where only a mixed life cycle is evolutionarily optimal. On both panels, we use rescaled variable for x-axis, 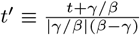.

Releasing the size constraints, we found that in near neutral environments, the evolutionary optimality of life cycles is independent on the seasons turnover period *T*. Life cycles evolve similarly in both the short and the long season regime, see Fig. 6A and Appendix K for a proof. As a consequence, in near neutral environments, the optimal pure life cycles can be inferred from the short seasons approximation. Since the areas of optimality map are separated by narrow borders in the order of ε in the short seasons regime, the pure binary fragmentation modes are evolutionarily optimal for the majority of dynamic environments with any season turnover period.

**Figure 6:**
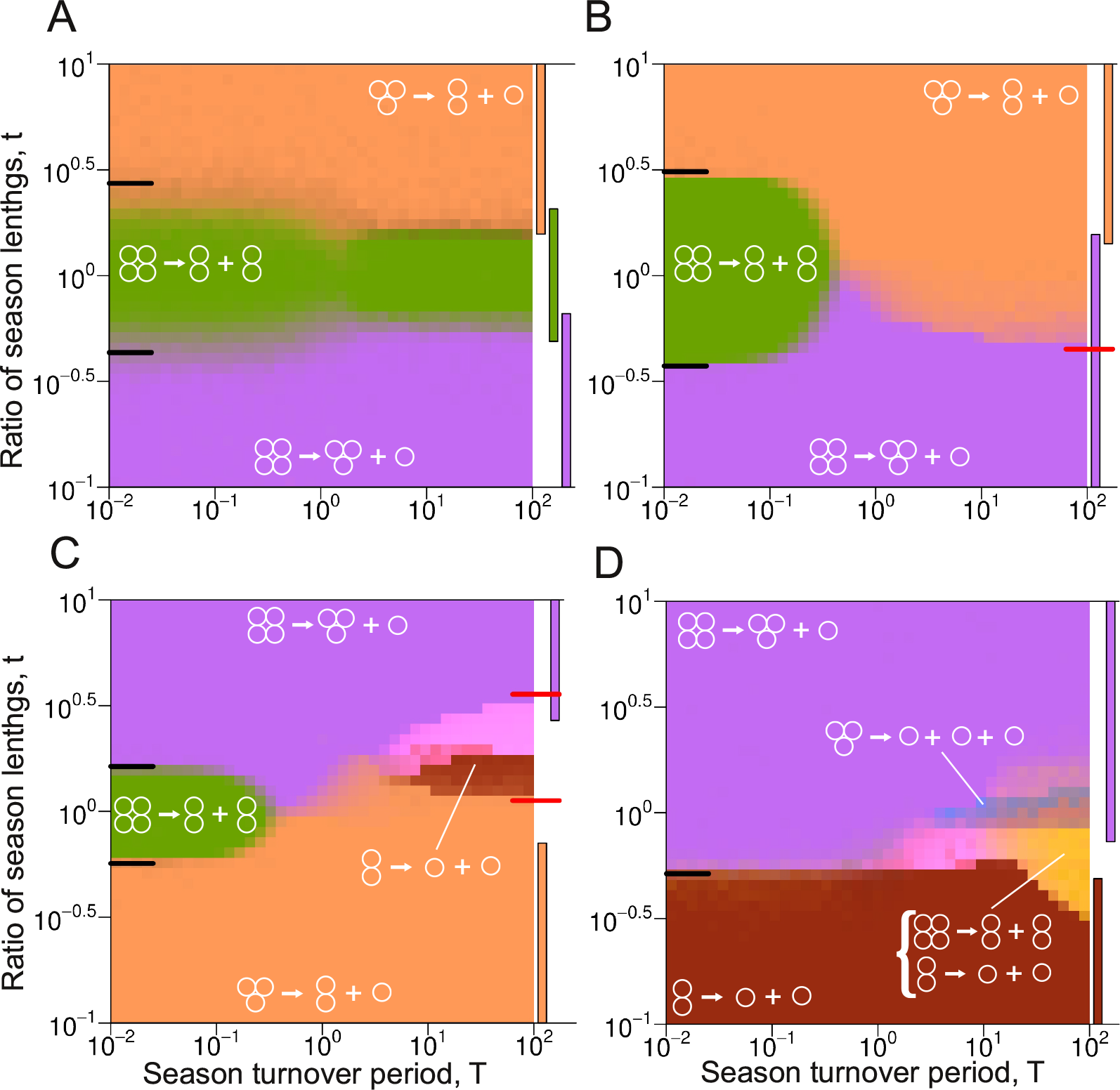
In the near-neutral dynamic environments, the long and the short seasons regimes promote similar life cycles, while in far-from-neutral environments, these can be very different. **A**. In the near neutral environment, the behaviour in the short and long seasons regime are similar. For environments far from neutral, this pattern is violated. **B**. A life cycle present in the short seasons regime (2+2) can be absent from the long seasons regime. **C**. Example in which a fragmentation pattern absent at the short seasons regime (1+1) contribute to the optimal life cycle in the long seasons regime. **D**. Example of a multiple fragmentation pattern (1+1+1, light blue) contributing to the optimal life cycle in the long seasons regime. Black lines on the left hand side of maps indicate borders predicted by the short seasons approximation, coloured bars to the right from maps indicate the locally stable areas of pure life cycles in the long seasons approximation. Red ticks on the right hand side of maps indicate the saddle-node bifurcations *t^∗^* in the long seasons approximation. Exact parameters of all presented calculation are presented in Appendix L.

If the birth rates are not in the vicinity of the neutral point *b*_*i*_ = 1, the set of optimal life cycles violates this scheme. Beyond the near neutral environment, we find more complex life cycles. We find that the fragmentation pattern presented in the short seasons regime may be absent in the long seasons regime, see Fig. 6B. The opposite is also possible, a fragmentation pattern absent in the short seasons may appear in the long seasons (as a component of a mixed life cycle, though), see Fig. 6C. Moreover, the multiple fragmentation, which cannot be evolutionarily optimal in any static environment, can evolve in a dynamic environment (again, as a component of the mixed life cycle), see Fig. 6D.

### 3.5 Robustness of pure binary fragmentation

While the potential diversity of the life cycles in dynamic environment is huge, an exotic behaviour is rare, and requires a fine balance of cell birth rates profiles {𝑆_1_, 𝑆_2_} and seasons lengths {τ_1_, τ_2_}. In the data set used in section 3.2, for each of 40, 000 pairs of seasons, we screened 41 different season length ratios (in total 1, 640, 000 dynamic environments), see Appendix E for details. For each environment, we found and characterized the set of evolutionarily optimal life cycles, see Fig. 7. The majority of dynamic environments (87%) promoted a unique pure life cycle. A smaller fraction (13%) featured coexistence of multiple local optima. A much smaller fraction of dynamic environments (1.9%) exhibited mixed life cycles, the majority of which was composed of two fragmentation patterns. Finally, multiple fragmentation was observed in a tiny set of environments (0.04%). Therefore, we conclude that pure life cycles should be a widespread evolutionary strategy in changing environment.

**Figure 7:**
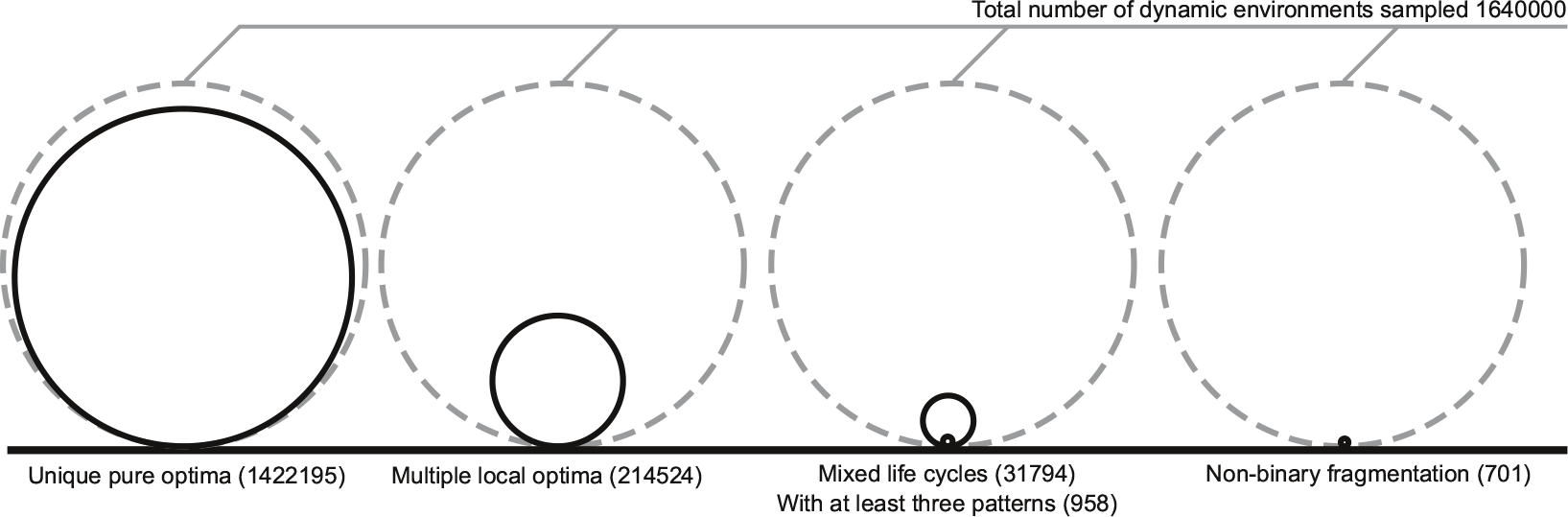
The majority of dynamic environments promotes a unique evolutionary optimal pure life cycle in a form of binary fragmentation. We sampled 1.64 · 10^6^ different long seasons dynamic environments, see Appendix E. More than one locally optimal fragmentation mode was found only in 13% of them (second circle). In 1.9% of dynamic environments, a set of locally optimal life cycles contained a mixed life cycle (third circle). Majority (97%) of found mixed life cycles executed just two fragmentation patterns. Only 3% of evolutionarily optimal mixed life cycles had three patterns or more. Finally, the number of dynamic environments promoted the evolution of non-binary fragmentation (1+1+1, 2+1+1, or 1+1+1+1) was extremely tiny - 0.04% (last circle). So, the most common result of life cycle optimization was a single local optimum, which happened to be a pure life cycle with binary fragmentation (first circle).

To support our result, we investigate the evolution of life cycles of larger colonial organisms. We consider groups growing up to size *n* = 15, so fragmentation must happen upon the birth of 16-th cell in a group. This size limit is comparable to the size of some volvocales algae, such as *Gonium pectorale* - one of the model organisms used to study the evolution of multicellularity.

We use the cell birth rate profiles 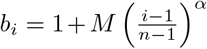. Investigation of these profiles in static environments [Pichugin et al., 2017] revealed that α ≪ 1 promotes an equal split (8 + 8), and α ≫ 1 promotes production of unicellular propagules (15 + 1). Note that the value of M has a relatively small influence. With these profiles, any increase in size is always beneficial to the group. Therefore, we restricted the optimization of life cycles to only fragmentation patterns of 16-cell groups. We obtain the optimality map, and the result supports our conclusion, see Fig. 8. Out of 1681 dynamic environments investigated, 1673 (99.5%) promoted a single unique optimal life cycle in a form of binary fragmentation. Only 8 environments (0.5%) had a mixed optimal life cycle, and no environment exhibited a coexistence of several local optima, or fragmentation into multiple pieces.

**Figure 8:**
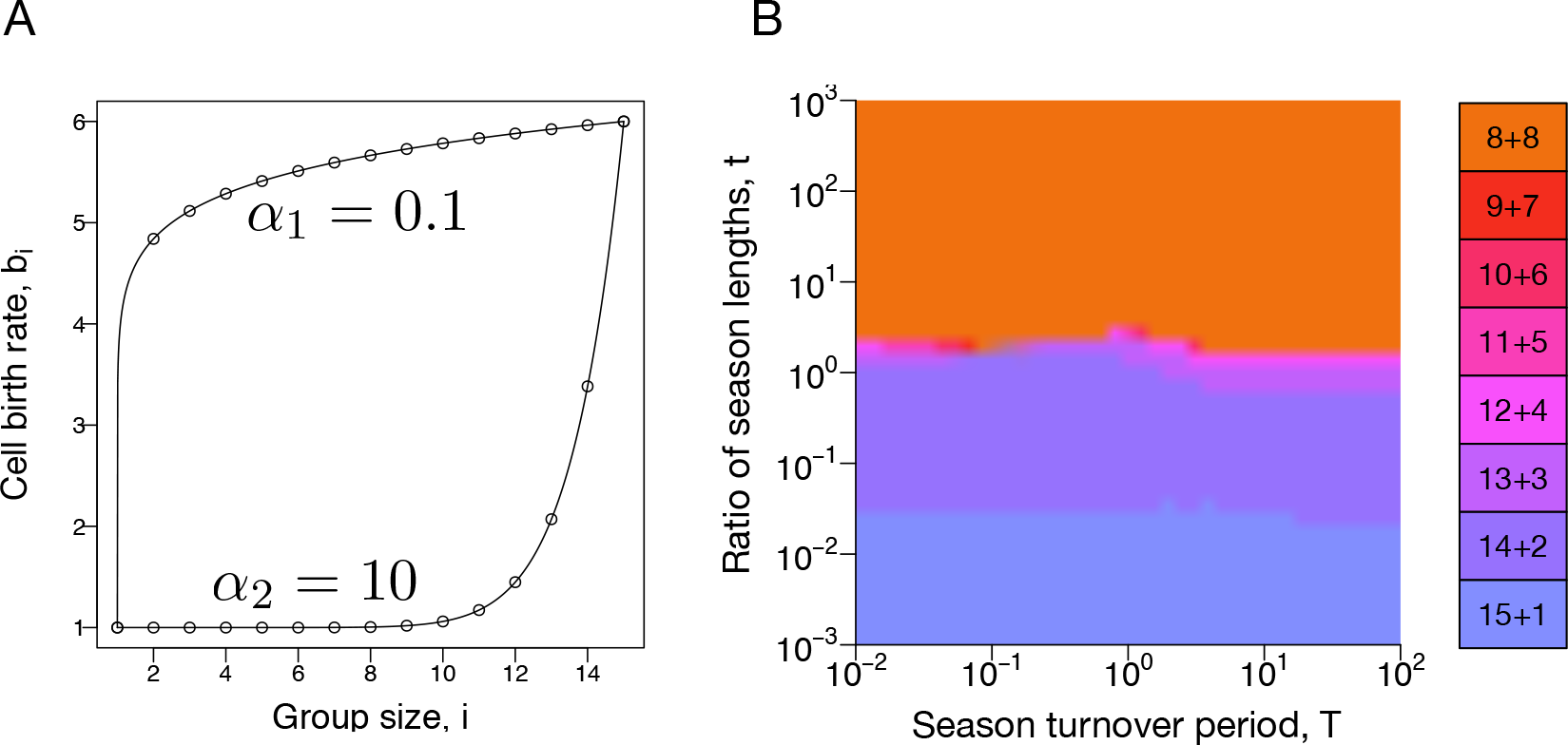
Monotonic dynamic environments promote binary fragmentation. **A**. To investigate the evolution of larger life cycles, we constructed a monotonic dynamic environment, where the birth rates *b*_*i*_ follow the function 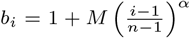 with different exponents *α* at different seasons. **B**. The majority of dynamic environments promoted unique pure life cycle with binary fragmentation. Even the environment is far from neutral, short and long seasons regimes exhibit similar evolutionary optimal life cycles. In our investigation, we used *M* = 5 in both seasons, *α* = 0.1 for the first season, and *α* = 10 for the second season. Due to significant computational load, we performed only 20 independent optimizations at each pixel of the map.

## 4 Discussion

Environmental fluctuations are commonly observed in nature. Under changing conditions, a trait beneficial in one season may become detrimental in another. Thus, adaptation to a dynamic environment may lead to totally different phenotypes than those that evolve in a static environment. In this manuscript, we investigated the influence of a changing environment on the evolution of life cycles in the context of primitive multicellularity. In our model, unstructured groups grow and eventually reproduce by fragmentation. The growth competition between different reproduction modes determines which life cycle will spread in population.

Our present model uses the minimal set of processes necessary for the multicellular life cycle: birth, growth, and reproduction of cell colonies. A number of other factors might influence the evolution of life cycles as well: aggregation of cells [Garcia and De Monte, 2013, Amado et al., 2018], group death [Pichugin et al., 2017], cell death [Amado et al., 2018, Pichugin and Traulsen, 2018], interactions between different cell types [Garcia et al., 2014, 2015, Gao et al., 2019], the geometry of groups [Libby et al., 2014], and so forth. However, given the current state of the field, the understanding of evolutionary dynamics of life cycles even in the minimal setups is missing. Our study has shown that even with basic processes, the spectrum of evolutionary outcomes is rich and deserves a dedicated investigation.

Previous findings reveal that a static environment puts strong constraints on evolutionarily optimal life cycles [Pichugin et al., 2017]. There, only pure life cycles can evolve. Moreover, among these, only binary fragmentation life cycles, featuring fragmentation into two groups, can become evolutionarily optimal. Interestingly, we found that evolution in dynamic environments can release both constraints.

Not only pure, but also mixed life cycles are able to evolve in our model. Being unable to perform well during both seasons, groups may employ a stochastic life cycle, where different groups randomly execute different fragmentation patterns. Thus, mixed life cycles are manifestation of between-clutch bet-hedging within the scope of our model. We found that in some dynamic environments, a mixed life cycle is the only evolutionarily optimal strategy, see Fig 5B. Our model also predicts that mixed life cycles may employ fragmentation patterns which would not contribute to the optimal life cycles under any of static seasonal components alone, see Fig. 6C and D. Yet, the most abundant scenario was the existence of only one locally optimal life cycle, which utilize a pure binary fragmentation mode, see Fig. 7. In other words, our model predicts that for simple multicellularity, between-clutch bet-hedging is possible to evolve, but is rarely an evolutionarily optimal strategy – even in changing environments.

Another form of bet-hedging observed among complex multicellular organisms is within-clutch bet-hedging, where offspring with diverse properties are produced in a single act of reproduction [Einum and Fleming, 2004]. From the perspective of simple multicellular life cycles, an act of reproduction is the distribution of the parental biomass among offspring. Hence, it is impossible to distinguish between an organism releasing a propagule and an organism producing two offspring of different size. In both cases, the result of reproduction is a collection of organisms of different sizes. The traditional point of view on these events is to consider them as asymmetric division, or propagule formation, and not as a bet-hedging scenario. In other words, while we can distinguish, who is the parent and who is the offspring in the case of chicken laying eggs, it is hardly possible to do so for a broken cyanobacteria filament. Thus, the very idea of within-clutch bet-hedging implies more developed multicellularity than one considered in our study.

In the light of the evolution of simple multicellularity, pure life cycles deserve special attention. One of the conceptual barriers for the species transitioning from unicellular to multicellular existence is the necessity to develop a predictable life cycle. While, the formal grouping of cells into clusters can give some advantages to the population [Rainey and Travisano, 1998, Ratcliff et al., 2012], the real strength of multicellularity eventually comes from beneficial interactions within the groups, such as cooperation or division of labour [Bell and Koufopanou, 1991, Kirk, 1997, Flores and Herrero, 2010, Hammerschmidt et al., 2014]. The ability of cells to participate in such interactions is not guaranteed beforehand and therefore, must evolve. For this, a regular schedule of a group growth and reproduction provides a proper basis. In our model, this regularity is obtained by pure life cycles. Thus, it is an interesting question, to which extent the changing environment can violate the evolutionary stability of pure life cycles.

Despite the mixed life cycles observed in simulations, pure life cycles remain prevalent: less than 2% of our dynamic environments promoted mixed cycles, see Fig. 7. Among the four analysed limiting regimes, three favour pure life cycles. If one season occupies a large enough proportion of the seasonal cycle, the evolutionary optimal life cycle is the same as if the second season does not happen at all. In other words, short disruptions of environmental conditions are unable to affect the evolutionary optimality of life cycles. The actual threshold below which the short season is unable to influence the life cycle evolution is a complex function of the environments and the turnover rate. Nevertheless, this value can be either inferred from numerical simulations of our model, or estimated from approximations. The short seasons regime only promotes pure life cycles. In near neutral environments, the behaviour at any seasons turnover time becomes similar to the one expressed at short seasons – the transitional area between pure life cycles become narrow. Only the long seasons regime can explicitly promote mixed life cycles. However, these emerge only at intermediary values of season length.

These findings are surprising because in dynamic environments, intuitively, mixed life cycles, which combine the best of both worlds, are expected to be optimal. Countering that intuition, we found that pure life cycles emerge for a wide range of dynamic environments.

