## Appendix for "Evolution of simple multicellular life cycles in dynamic environments"

#### A Equation for the population dynamics in a matrix form

In our model, groups grow in size and fragment. The growth of groups is governed by a set of biological reactions

$$X_i \xrightarrow{ib_i g_i} X_{i+1}, \tag{13}$$

where $X_i$ is a groups of size $i$ and $b_i$ is the growth rate of cells in a group of size $i$. After the growth of the group size by $i+1$, they will stay together with the probability $g_i$, which is determined by the utilized life cycle and the group size.

Group fragmentation occurs immediately after cell growth and is described by reactions in the form

$$X_i \xrightarrow{ib_i q_\kappa} \sum_{j=1}^{i} \pi_j(\kappa) X_j, \tag{14}$$

where $\kappa$ indicates the pattern of the fragmentation (for example 2+1+1), and $\pi_j(\kappa)$ indicates how many groups of size $j$ are produced by fragmentation $\kappa$ (e.g. if $\kappa = 2+1+1$, then $\pi_1(\kappa) = 2$ and $\pi_2(\kappa) = 1$). The probabilities $q_\kappa$ determine the fragmentation mode of a given life cycle.

At each size of group multiple patterns of fragmentation are available. For instance, upon reaching size of 3 cells, a group may fragment into a two-cellular group and independent cell (pattern 2+1), or into three solitary cells (pattern 1+1+1). To denote that a group of the size $i$ can fragment according to the pattern $\kappa$, we write $\kappa \vdash i$, so in previous example $2+1 \vdash 3$ and $1+1+1 \vdash 3$. At each size, the sum of probabilities to stay together or to fragment is equal to one

$$g_i + \sum_{\kappa \vdash i+1} q(\kappa) = 1. \tag{15}$$

The sets of reactions (13) and (14) give rise to a system of differential equations

$$\dot{x_i} = \sum_{j=1}^{n} \sum_{\kappa \vdash j+1} q_\kappa \pi_i(\kappa) j b_j x_j - i b_i x_i + g_{i-1}(i-1) b_{i-1} x_{i-1}, \tag{16}$$

where $x_i$ denotes the abundance of groups of size $i$ and $n$ is the maximal size of groups present in population. In this study we use $n = 3$. The equations (16) are linear and can be represented in matrix form,

$$\dot{\mathbf{x}} = A\mathbf{x}, \tag{17}$$

where $\mathbf{x} = (x_1, x_2, x_3, \cdots)$ is the vector of group size abundances, and $A$ is the projection matrix with elements are given by

$$A_{i,j}(\mathbf{q}, b_j) = jb_j \left( \sum_{\kappa \vdash j+1} q_\kappa \pi_i(\kappa) - \delta_{i,j} + g_j \delta_{i,j+1} \right). \tag{18}$$

### B Numerical calculation of the growth rate in a dynamic environment

The growth rate of a population in a dynamic environment is calculated as a slope of a linear fit of the logarithm of the population size vs. time. Each simulation of a growing populations begins with a population of a random composition, where the abundances of each group size (solitary cells, 2- and 3-cellular groups) are drawn independently from a uniform distribution $U(0, 1)$. Using random initial states, we were able to explicitly measure the impact of stochasticity in the initial conditions on the calculated value of the population growth rate (see below). Also, since our initial state include multicellular complexes, we could correctly handle the population dynamics of the coexisting life cycle.

For every season of the dynamic environment, the eigenvalues $(\lambda_i)$ and eigenvectors $(\mathbf{e}_i)$ of the projection matrix $A$ are computed. After that, the vector of the initial composition of the population was decomposed into the basis of the eigenvectors. Eventually, each component of such decomposition exponentially grows in time independently with a rate given by the eigenvalue associated with the eigenvector. This allows us obtain the population composition at the end of the season, or at any moment during the season as

$$\mathbf{x}(\tilde{t}) = \sum_i c_i(0) \mathbf{e}_i e^{\lambda_i \tilde{t}}, \tag{19}$$

where $\tilde{t}$ is time, $c_i(0)$ is the weight of the eigenvector $\mathbf{e}_i$ in the initial state of population $\mathbf{x}(0)$, and the sum runs over all eigenvalues.

The population composition achieved at the end of the first season is used as an initial composition at the beginning of the second season (where eigenvalues and eigenvectors of the projection matrix are totally different). To calculate the growth rate in the dynamic environment $(\Lambda)$, we apply two alternating seasons for a long time and compute population sizes. The best linear fit of the logarithm of population size gives the growth rate $\Lambda$ of population.

The random initial conditions are the source of variation in $\Lambda$ between independent calculations with identical parameters ($\mathcal{D}$, $\mathbf{q}$, total simulation time). The longer is the total simulation time, the less is the variation in $\Lambda$ caused by the initial conditions. The question is, how long this should be? For the purpose of our study, $\Lambda$ must be obtained with an accuracy allowing a reliable comparison of growth rates with differences in fragmentation probabilities of 0.05. This value is the size of the lattice on which the optimization is performed, see appendix C.

Preliminary simulations show that such a difference in fragmentation probabilities induce a relative difference between growth rates of the order of $10^{-2}$. The next set of preliminary simulations show that to achieve such an accuracy, the total simulation time should last for at least 10 units of time ($\tilde{t}$). This (very roughly) corresponds to 10 generations in a population utilizing the unicellular life cycle 1+1. In our simulations, we used the limit of 30 time units. Furthermore, we investigate regimes, where the cycle of seasons was longer than 30 time units (so called long seasons regime). For these regimes, simulations must run longer, to accomodate at least several seasons turnovers. Therefore, to adequately compute $\Lambda$ in these cases as well, we also require our calculations to last for minimum 20 full cycles of seasons change.

### C Numerical optimization of the growth rate

For a given dynamic environment $\mathcal{D}$, we find the local optima of the function $\Lambda(\mathbf{q}, \mathcal{D})$ varying $\mathbf{q} = (q_{1+1}; q_{2+1}, q_{1+1+1}; q_{3+1}, q_{2+2}, q_{2+1+1}, q_{1+1+1+1})$. The set of fragmentation probabilities for a maximum size of 4 has to satisfy the conditions $q_{1+1} \leq 1$, $q_{2+1}+q_{1+1+1} \leq 1$ and $q_{3+1}+q_{2+2}+q_{2+1+1}+q_{1+1+1+1} = 1$. To investigate this six-dimensional space of parameters, we set up a lattice with spacing 0.05. The value of $\Lambda(\mathbf{q}, \mathcal{D})$ is computed for all nodes of this lattice. The whole lattice contains $\frac{21^3 \cdot 22^2 \cdot 23}{12} \approx 8.6 \cdot 10^6$ nodes, and the computation $\Lambda$ in each of them would be inefficient. Thus, we implemented a hill climbing optimization algorithm. The optimization begins from a random node. Then, the values of $\Lambda$ in the neighboring nodes are calculated, and the node with largest $\Lambda$ is chosen for the next step. Once, there is no neighboring node with a larger $\Lambda$, the vector $\mathbf{q}$ is considered as a candidate to the local optimum. Each single optimization procedure returns a single local optimum. However, multiple local optima may exist. To capture them, we repeat the optimization procedure 100 times with random initial values of $\mathbf{q}$.

Note that if the fragmentation always happens at a group size $l$ smaller than the maximum group size $n$, the fragmentation probabilities $q_\kappa$ at larger group sizes $l' > l$ do not affect $\Lambda$. For $l = 4$, the single exceptional case is the coexisting life cycle $q_{1+1} = 1$ and $q_{2+1} + q_{1+1+1} = 0$ and $q_{2+2} = 1$. However, while the value of $\Lambda$ is independent on fragmentation probabilities at unavailable sizes, the gradient of the growth rate can still depend on them. For instance, at $\mathcal{S} = (1, 4, 2)$, the parameter combination $\mathbf{q}_1 = (0; 1, 0; 0.1, 0.3, 0.55, 0.05)$ corresponds to the pure life cycle 2+1 and does not have a neighboring nodes with larger $\Lambda$. However, the lattice node $\mathbf{q}_2 = (0; 1, 0; 0, 1, 0, 0)$ corresponds to the same pure life cycle 2+1 but has a neighboring node with larger $\Lambda$, so the optimization procedure can further improve the growth rate. To handle this issue we added the second round of optimization. If the originally found candidate for the local optimum satisfies $q_{2+1} + q_{1+1+1} = 1$ or $q_{1+1} = 1$, we initialize the set of optimizations starting from the same life cycle but with probability mass of unused partitions altered to be concentrated in each of these partitions. So, in the example above, after the hill climber algorithm reports the above-mentioned $\mathbf{q}_1$ as being the candidate to the local optimum, we set up four additional instances of optimization starting from:

- $\mathbf{q}_a = (0; 1, 0; |1, 0, 0, 0)$
- $\mathbf{q}_b = (0; 1, 0; |0, 1, 0, 0)$
- $\mathbf{q}_c = (0; 1, 0; |0, 0, 1, 0)$
- $\mathbf{q}_d = (0; 1, 0; |0, 0, 0, 1)$

where we separated probabilities of fragmentation at size 4 by a vertical line for the convenience of presentation.

Once the local optimum in a given simulation is found, we clear the fragmentation mode by removing unused fragmentation probabilities. We take into account the following cases:

- If $q_{1+1} = 1$, then groups of size larger than 1 do not emerge, so all other probabilities can be discarded independently on their values, except the coexisting life cycle $\mathbf{q}_C$ (see main text).
- If $q_{2+1} + q_{1+1+1} = 1$, then groups of size larger than 2 do not emerge, such that the probabilities $q_{3+1}, q_{2+2}, q_{2+1+1}, q_{1+1+1+1}$ can be discarded independently on their values.
- If $q_{2+1} + q_{1+1+1} = 0$ and $q_{2+2} = 1$, then groups of size 1 do not emerge, so the probability $q_{1+1}$ can be discarded independently on its value, except the coexisting life cycle.
- If $q_{1+1} = 1$ and $q_{2+1} + q_{1+1+1} = 0$ and $q_{2+2} = 1$, then both 1+1 and 2+2 life cycles are executed simultaneously, since these are coexisting life cycles. This case is kept intact.

### D Colour code of the optimality maps

The optimality maps represent the character of the evolutionarily optimal life cycle(s) as a function of the control parameters $T$ and $t$. Each pixel of the map corresponds to a single dynamic environment $\mathcal{D} = \{\mathcal{S}_1, \tau_1; \mathcal{S}_2, \tau_2\}$ and its colour represents the set of evolutionary optimal life cycles found in the given environment. Each optimization results in a single locally optimal fragmentation mode $\mathbf{q}$. For each dynamic environment, we performed 100 such runs leading to a list of 100 local optima (containing repeated entries). We denote the probability of the fragmentaiton pattern $\kappa$ to be executed in the evolutionarily optimal life cycle found in $i$-th optimization as $q_{\kappa,i}$. Here, we describe how we mapped a list of $q_{\kappa,i}$ into the colour of pixel on the optimality map.

We started with our fragmentation pattern colour code introduced in Fig. 1. The list of colours was generated to have the maximal perception distance between seven colours using web tool http://tools.medialab.sciences-po.fr/iwanthue/. Colours we used are:

- Pattern 1+1, hex code #992b10, dark red
- Pattern 2+1, hex code #ff9a58, light orange
- Pattern 1+1+1, hex code #00abfd, blue
- Pattern 3+1, hex code #c46bf4, soft violet
- Pattern 2+2, hex code #6ba400, dark green
- Pattern 2+1+1, hex code #cd5a87, moderate pink
- Pattern 1+1+1+1, hex code #b3713f, brown

Then we compute the average occurrence of each fragmentation pattern across all found optima as $\langle q_\kappa \rangle = \frac{1}{100}\sum_i q_{\kappa,i}$, where $\kappa$ stands for a fragmentation pattern and summation goes over all 100 entries of the local optima list.

After that, we convert the found "average" life cycle into the pixel colour. To construct the colour of a mixed life cycle $\mathbf{q} = (q_{1+1}; q_{2+1}, q_{1+1+1}; q_{3+1}, q_{2+2}, q_{2+1+1}, q_{1+1+1+1})$, we evaluate the weighted sum of these base colours in RGB space. The weight of each colour $w_\kappa$ is equal to the product of the corresponding pattern frequency by the probability that a group of size one cell smaller will grow and not fragment:

$$w_\kappa = q_\kappa \left(1 - \sum_{\kappa' \vdash s-1} q_{\kappa'}\right) \tag{20}$$

This weighting procedure provides an illustration of actual frequencies of fragmentation pattern occurrences. Consider a mixed life cycle $\mathbf{q} = (0.95; 0, 1; 0, 0, 0, 0)$, where 95% of independent cells immediately fragment upon growth and the remaining 5% form bi-cellular cluster, which fragment upon the next division as 1+1+1. The frequency of this pattern is $q_{1+1+1} = 1$, however, it is responsible for the minority of reproduction events in the population. Our colour mixing rule assigns a weight of the colour corresponding to $\kappa = 1+1+1$ equal to $q_{1+1+1}(1 - q_{1+1}) = 0.05$, which is more appropriate in this situation.

### E Long seasons dynamical environment screening

Evolution of fragmentation modes in the long seasons regime can lead to many different outcomes. To investigate the spectrum of evolutionarily optimal life cycles and their combinations, we performed screening in a wide range of dynamic environments. In this screening, the parameters of the first season $\mathcal{S}_1 = (1, b_2^1, b_3^1)$ were sampled from the set $\{0.25, 0.75, 1.25, \ldots, 4.75\}$, ten values in total. Altogether, this created 100 different $\mathcal{S}_1$. The parameters of the second season $\mathcal{S}_2 = (1, b_2^2, b_3^2)$ were independently sampled from a uniform distribution $U(0, 5)$. For each $\mathcal{S}_1$, 400 different $\mathcal{S}_2$ were generated. For each combination of $\mathcal{S}_1$ and $\mathcal{S}_2$, 41 equally spaced values of $\log(t)$ in the range $[-1, 1]$ were assessed numerically. In total, we investigated $100 \times 400 \times 41 \approx 1.6 \cdot 10^6$ dynamic environments. Out of these, the vast majority contained only a single evolutionarily optimal pure life cycle. A coexistence between several local optima has been observed in approximately 13% of dynamic environments, see Table 1. Mixed life cycles were found among local optima in 1.9% of cases, and only in 0.06% of considered dynamic environments, these executed more than two fragmentation patterns. Multiple fragmentation patterns (1+1+1, 2+1+1, or 1+1+1+1) were found to be even rarer, 0.04%.

| | Single pure life cycle | Multiple local optima | Mixed life cycle | Mixed life cycles with 3+ patterns | Non-binary fragmentation | Total |
|---|---|---|---|---|---|---|
| Counts | 1422195 | 214524 | 31794 | 958 | 701 | 1640000 |
| Fraction | 0.867 | 0.131 | 0.0193 | 0.000584 | 0.000427 | 1.0 |

Table 1: The majority of evolutionarily optimal life cycles in the long seasons regime are single pure life cycles.

### F Stability of pure life cycles in the long seasons regime

Here we consider evolutionary optimality of pure life cycles, where only one fragmentation pattern occurs. A pure fragmentation mode is locally optimal when adding a small chance to fragment with any other pattern leads to a decrease in the population growth rate $\Lambda$. The stability analysis of a pure life cycle can then be reduced to a set of pairwise comparisons.

In each comparison, we examine the stability of the focal pure fragmentation mode $\mathbf{q}_f$ against the perturbation in the direction of the alternative pure fragmentation mode $\mathbf{q}_p$. To do this, we consider a mixed life cycle executing only two fragmentation patterns: the focal $\kappa_f$ and the perturbation $\kappa_p$. Such a life cycle is characterized by the fragmentation mode $\mathbf{q}_{f,p}(x)$, where patterns $\kappa_f$ and $\kappa_p$ occur with probabilities $x$ and $1-x$, respectively. The pure focal fragmentation mode $\mathbf{q}_f$ is obtained at $x = 1$ and is locally stable with respect to admixture of $\mathbf{q}_p$ if the growth rate is increasing near $x = 1$, $\frac{\partial \Lambda(\mathbf{q}_{f,p}(x), \mathcal{D})}{\partial x}|_{x=1} \equiv \Lambda'_x|_{x=1} > 0$.

To find the ratio of season lengths $t_s$ at which the life cycle $\mathbf{q}_f$ becomes (un)stable against $\mathbf{q}_p$, consider the long seasons approximation:

$$\Lambda(\mathbf{q}_{f,p}, \mathcal{D}) \approx \frac{t}{1+t}\lambda(\mathbf{q}_{f,p}, \mathcal{S}_1) + \frac{1}{1+t}\lambda(\mathbf{q}_{f,p}, \mathcal{S}_2). \tag{21}$$

and therefore

$$\Lambda'_x(\mathbf{q}_{f,p}, \mathcal{D})|_{x=1} \approx \frac{t_s(\kappa_f, \kappa_p)}{1 + t_s(\kappa_f, \kappa_p)}\lambda'_x(\mathbf{q}_{f,p}, \mathcal{S}_1)|_{x=1} + \frac{1}{1 + t_s(\kappa_f, \kappa_p)}\lambda'_x(\mathbf{q}_{f,p}, \mathcal{S}_2)|_{x=1} = 0. \tag{22}$$

Solving this equation with respect to $t_s(\kappa_f, \kappa_p)$, we get

$$t_s(\kappa_f, \kappa_p) = -\left.\frac{\lambda'_x(\mathbf{q}_{f,p}(x), \mathcal{S}_2)}{\lambda'_x(\mathbf{q}_{f,p}(x), \mathcal{S}_1)}\right|_{x=1}. \tag{23}$$

Note that Eq. (23) links properties of a pure life cycle in a dynamic environment ($t_s$) with growth rates of mixed life cycle in a static environment ($\lambda'_x$).

If both seasons favours $\mathbf{q}_f$ over $\mathbf{q}_p$, then $t_s < 0$, and $\mathbf{q}_f$ is locally stable for any $t$. If both seasons favours $\mathbf{q}_p$ over $\mathbf{q}_f$, then $t_s < 0$, and $\mathbf{q}_f$ is locally unstable for any $t$. If the first season favours $\mathbf{q}_f$ over $\mathbf{q}_p$, and the second season is opposite $[\lambda(\mathbf{q}_f, \mathcal{S}_1) > \lambda(\mathbf{q}_p, \mathcal{S}_1)$ and $\lambda(\mathbf{q}_f, \mathcal{S}_2) < \lambda(\mathbf{q}_p, \mathcal{S}_2)]$, then $\mathbf{q}_f$ is locally stable at $t > t_s$. Finally, if the relation is inverse $[\lambda(\mathbf{q}_f, \mathcal{S}_1) < \lambda(\mathbf{q}_p, \mathcal{S}_1)$ and $\lambda(\mathbf{q}_f, \mathcal{S}_2) > \lambda(\mathbf{q}_p, \mathcal{S}_2)]$, then $\mathbf{q}_f$ is locally stable at $t < t_s$.

The pure fragmentation mode $\mathbf{q}_f$ is locally optimal if it is locally stable against all other pure fragmentation modes $\mathbf{q}_{p \neq f}$. Therefore, the range of $t$ where $\mathbf{q}_f$ is locally optimal is given by the intersection of stability regions obtained from each pairwise assessment.

### G Stability of mixed life cycles in the long seasons regime

In this section, we outline the range of season length ratios $t$ promoting mixed life cycles. An arbitrary mixed fragmentation mode $(\mathbf{q})$ is a local optimum of the growth rate $\Lambda$, when all fragmentation patterns $\kappa$ fulfils the conditions

$$\begin{cases} \left.\frac{\partial \Lambda}{\partial q_\kappa}\right|_{\mathbf{q}} < 0 & \text{if } \kappa \text{ is not executed } (\mathbf{q}_\kappa = 0), \\ \left.\frac{\partial \Lambda}{\partial q_\kappa}\right|_{\mathbf{q}} = 0 \text{ and } \left.\frac{\partial^2 \Lambda}{\partial q_\kappa^2}\right|_{\mathbf{q}} < 0 & \text{if } \kappa \text{ is mixed with other patterns of the same size } (0 < \mathbf{q}_\kappa < 1), \\ \left.\frac{\partial \Lambda}{\partial q_\kappa}\right|_{\mathbf{q}} > 0 & \text{if } \kappa \text{ is the only executed pattern of its group size } (\mathbf{q}_\kappa = 1). \end{cases} \tag{24}$$

The majority of mixed life cycles we found features only two fragmentation patterns, see Appendix H. For this case, the analysis can be significantly simplified, and we can explicitly find the range of $t$, where the mixed fragmentation modes are evolutionarily optimal. Let us consider again a pair of fragmentation patterns corresponding to pure fragmentation modes $\mathbf{q}_1$ and $\mathbf{q}_2$, such as $\mathcal{S}_1$ favours $\mathbf{q}_1$ and $\mathcal{S}_2$ favours $\mathbf{q}_2$ $[\lambda(\mathbf{q}_1, \mathcal{S}_1) > \lambda(\mathbf{q}_2, \mathcal{S}_1)$ and $\lambda(\mathbf{q}_1, \mathcal{S}_2) < \lambda(\mathbf{q}_2, \mathcal{S}_2)]$. Now we focus on the behaviour of the locally optimal mixed fragmentation mode $\mathbf{q}_{1,2}(x_m)$ with $\Lambda'_x|_{x=x_m} = 0$ and $\Lambda''_{xx}|_{x=x_m} \leq 0$, where $x_m \neq 1, 0$.

Above, we have shown that there are no evolutionarily optimal mixed life cycles if there is effectively a single season ($t \ll 1$ or $t \gg 1$). Therefore, as approaching these extreme values of $t$, $x_m$ either hits 0 or 1 (see Fig. 1A) or disappears in a saddle-node bifurcation (see Fig. 1B).

In the first scenario, if $x_m$ converges to 1 as $t$ increases, the internal maximum becomes the border maximum (evolutionarily stable pure life cycle). This happens at $t = t_s(\kappa_1, \kappa_2)$ by the definition of $t_s$. However, the inverse is not always true; not every $t_s$ marks the transition between the border and the internal maxima, because the same condition is satisfied when the border maximum merges with an internal minimum. For an emergence of internal maximum, $\Lambda''_{xx}|_{x=1} \leq 0$ must be satisfied at $t = t_s$, i.e.

$$\Lambda''_{xx}(\mathbf{q}_{1,2}(x), \mathcal{D})|_{x=1} \approx \frac{t_s(\kappa_f, \kappa_p)}{1 + t_s(\kappa_f, \kappa_p)} \lambda''_{xx}(\mathbf{q}_{1,2}(x), \mathcal{S}_1)|_{x=1} + \frac{1}{1 + t_s(\kappa_f, \kappa_p)} \lambda''_{xx}(\mathbf{q}_{1,2}(x), \mathcal{S}_2)|_{x=1} < 0. \tag{25}$$

Using Eq. (23) and rearranging terms, we get

$$-\left.\left( \frac{\lambda'_x(\mathbf{q}_{1,2}(x), \mathcal{S}_2)}{\lambda'_x(\mathbf{q}_{1,2}(x), \mathcal{S}_1)} \lambda''_{xx}(\mathbf{q}_{1,2}(x), \mathcal{S}_1) \right)\right|_{x=1} + \lambda''_{xx}(\mathbf{q}_{1,2}(x), \mathcal{S}_2)|_{x=1} < 0. \tag{26}$$

If $\mathcal{S}_1$ promotes $\mathbf{q}_f$ over $\mathbf{q}_p$ while $\mathcal{S}_2$ promotes $\mathbf{q}_p$ over $\mathbf{q}_f$, then $\lambda'_x(\mathbf{q}_{1,2}(x), \mathcal{S}_2) < 0$, so the inequality can be rewritten as

$$\left.\frac{\lambda''_{xx}(\mathbf{q}_{1,2}(x), \mathcal{S}_1)}{\lambda'_x(\mathbf{q}_{1,2}(x), \mathcal{S}_1)}\right|_{x=1} < \left.\frac{\lambda''_{xx}(\mathbf{q}_{1,2}(x), \mathcal{S}_2)}{\lambda'_x(\mathbf{q}_{1,2}(x), \mathcal{S}_2)}\right|_{x=1}. \tag{27}$$

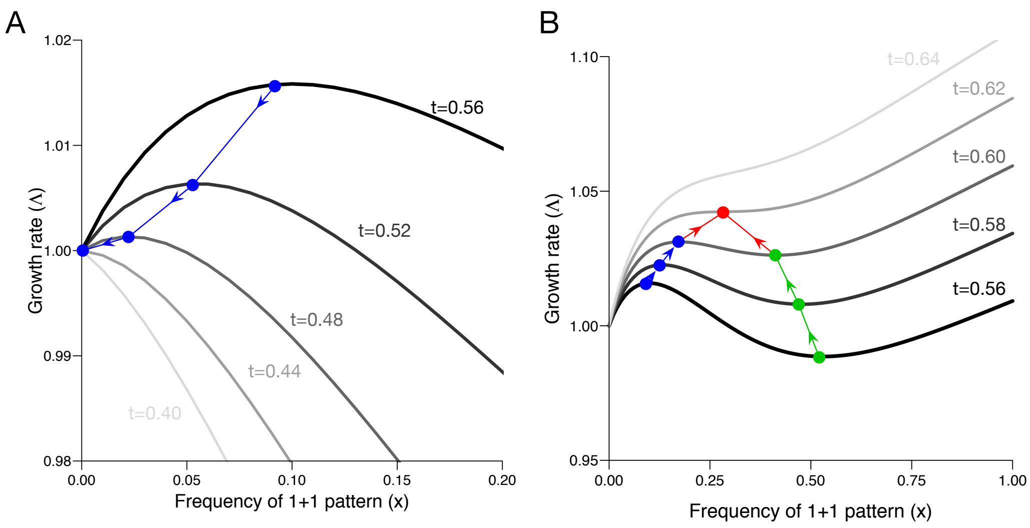


Figure 1: **At extreme $t$, locally optimal mixed life cycles either become pure or disappear in a saddle-node bifurcation.** Consider a mixed life cycle executing two patterns 1+1 and 2+1, i.e. $\mathbf{q} = (x; 1, 0; 0, 0, 0, 0)$. In a dynamic environment given by $\mathcal{S}_1$ = (1.0, 2.0, 0.1) and $\mathcal{S}_2$ = (1.0, 0.2, 0.1), at $T \gg 1$ and $t = 0.56$ (thick black lines on both panels), the maximal growth rate is obtained at $x_m \approx 0.10$ (highlighted with a blue dot). **A** With decrease in $t$, the location of the optimal mixed life cycle $x_m$ goes left, until it hits $x_m = 0$ at $t_s \approx 0.44$. There is no evolutionary optimal mixed life cycles at $t < t_s$. **B** With increase in $t$, the location of the optimal mixed life cycle $x_m$ goes right. At the same time, the location of local minimum of $\Lambda$ goes left (highlighted with green dots). At $t^* \approx 0.62$, local maximum disappears in the saddle-node bifurcation located at $x^* \approx 0.27$ (the red dot). There is no evolutionary optimal mixed life cycles at $t > t^*$.

If $\mathcal{S}_1$ promotes $\mathbf{q}_p$ instead, the inequality sign is reversed. For transitions at another boundary, the same expressions should be evaluated at $x = 0$, instead of $x = 1$.

In the scenario of the saddle-node bifurcation, the internal maximum and minimum of the growth rate merge, see Fig. 1B. Then, there is a critical values of $x = x^*$ and $t \equiv t^*$, at which the growth rate profile $\Lambda(\mathbf{q}(x), \mathcal{D})$ has an equilibrium which is simultaneously an inflection point:

$$\begin{aligned} \Lambda'_x(\mathbf{q}_{1,2}(x), \mathcal{D})\big|_{x=x^*} &= 0, \\ \Lambda''_{xx}(\mathbf{q}_{1,2}(x), \mathcal{D})\big|_{x=x^*} &= 0. \end{aligned} \tag{28}$$

Under the long seasons approximation, we get

$$\begin{aligned} t^* &= -\left.\frac{\lambda'_x(\mathbf{q}_{1,2}(x), \mathcal{S}_2)}{\lambda'_x(\mathbf{q}_{1,2}(x), \mathcal{S}_1)}\right|_{x=x^*}, \\ t^* &= -\left.\frac{\lambda''_{xx}(\mathbf{q}_{1,2}(x), \mathcal{S}_2)}{\lambda''_{xx}(\mathbf{q}_{1,2}(x), \mathcal{S}_1)}\right|_{x=x^*}. \end{aligned} \tag{29}$$

Setting the two equations equal and rearranging the terms, we get

$$\left.\frac{\lambda''_{xx}(\mathbf{q}_{1,2}(x), \mathcal{S}_1)}{\lambda'_x(\mathbf{q}_{1,2}(x), \mathcal{S}_1)}\right|_{x=x^*} = \left.\frac{\lambda''_{xx}(\mathbf{q}_{1,2}(x), \mathcal{S}_2)}{\lambda'_x(\mathbf{q}_{1,2}(x), \mathcal{S}_2)}\right|_{x=x^*}. \tag{30}$$

For the border of the second type to exists, there must be at least one $x^*$ satisfying Eq. (30). If none of these are found, such a dynamic environment promotes only pure life cycles.

### H Growth rate of a mixed life cycle constructed by two fragmentation patterns in a static environment

In this section, we present the calculation of the growth rates of the mixed life cycle utilizing exactly two different fragmentation patterns. This value is a solution of the characteristic equation of the projection matrix, which we aim to derive. We begin with the projection matrix of a pure life cycle (see [Pichugin et al., 2017] for details),

$$A = \begin{pmatrix} -b_1 & 0 & \cdots & 0 & (\ell-1)b_{\ell-1}\pi_1 \\ b_1 & -2b_2 & 0 & \vdots & (\ell-1)b_{\ell-1}\pi_2 \\ 0 & 2b_2 & -3b_3 & 0 & (\ell-1)b_{\ell-1}\pi_3 \\ 0 & 0 & \ddots & \ddots & \vdots \\ 0 & 0 & \cdots & (\ell-2)b_{\ell-2} & (\ell-1)b_{\ell-1}\left(\pi_{\ell-1}-1\right) \end{pmatrix},$$

where $b_i$ is the cell birth rate in a group size $i$, and $\ell$ is the size at which a fragmentation occurs, and $\pi_i$ is the number of groups with size $i$ produced in a result of the fragmentation. This matrix contains non-zero components only at the main diagonal, the lower sub-diagonal (growth components) and the rightmost $(\ell-1)$-th column (fragmentation components). To find the growth rate $\lambda$, the characteristic equation $\det(A - \lambda I) = 0$ must be solved. By dividing the $i$-th column of $A - \lambda I$ by $ib_i$, we get:

$$\det \begin{pmatrix} -s_1 & 0 & \cdots & 0 & \pi_1 \\ 1 & -s_2 & 0 & \vdots & \pi_2 \\ 0 & 1 & -s_3 & 0 & \pi_3 \\ 0 & 0 & \ddots & \ddots & \vdots \\ 0 & 0 & \cdots & 1 & -s_{\ell-1}+\pi_{\ell-1} \end{pmatrix} = 0, \tag{31}$$

where $s_i = \left(1 + \frac{\lambda}{ib_i}\right)$ – this notation will be used later for the convenience of the matrix presentation. The characteristic equation can be rewritten in a form of the characteristic polynomial:

$$p(\lambda) = F_\ell(\lambda) - \sum_{i=1}^{\ell-1} \pi_i F_i(\lambda) = 0, \tag{32}$$

where

$$F_i(\lambda) = \prod_{j=1}^{i-1} s_j = \prod_{j=1}^{i-1} \left(1 + \frac{\lambda}{jb_j}\right). \tag{33}$$

For instance, the growth rate of the pure life cycle 2+1 is the largest root of the polynomial:

$$p_{2+1}(\lambda) = F_3 - F_2 - F_1 = \left(1 + \frac{\lambda}{b_1}\right)\left(1 + \frac{\lambda}{2b_2}\right) - \left(1 + \frac{\lambda}{b_1}\right) - 1 = 0.$$

We define a mixed life cycle $\mathbf{q}(x)$, as a life cycle in which an initially single-cellular group would eventually fragment according to $\kappa_1$ with probability $x$ and according to $\kappa_2$ with probability $1-x$. If we denote the size at which fragmentation occurs in the fragmentation pattern $\kappa_i$ as $\ell_i$, then the fragmentation mode mixed between two pure modes $\mathbf{q}_1$ and $\mathbf{q}_2$ is defined as

$$\mathbf{q}(x) = \begin{cases} x\mathbf{q}_1 + \mathbf{q}_2, & \text{if } \ell_1 < \ell_2, \\ x\mathbf{q}_1 + (1-x)\mathbf{q}_2, & \text{if } \ell_1 = \ell_2, \\ \mathbf{q}_1 + (1-x)\mathbf{q}_2, & \text{if } \ell_1 > \ell_2. \end{cases} \tag{34}$$

At the boundary values of $x$, the mixed life cycle becomes pure $\mathbf{q}(0) = \mathbf{q}_2$ and $\mathbf{q}(1) = \mathbf{q}_1$ (a fragmentation at a larger size does not happen if all groups fragment at a smaller size). Thus, the characteristic equation is given by

$$\det \begin{pmatrix} -s_1 & 0 & \cdots & 0 & \pi_1(\kappa_1)x & 0 & \cdots & 0 & \pi_1(\kappa_2) \\ 1 & -s_2 & \cdots & 0 & \pi_2(\kappa_1)x & 0 & \cdots & 0 & \pi_2(\kappa_2) \\ \vdots & \vdots & \ddots & \vdots & \vdots & \vdots & \ddots & \vdots & \vdots \\ 0 & 0 & \cdots & -s_{\ell_1-2} & \pi_{\ell_1-2}(\kappa_1)x & 0 & \cdots & 0 & \pi_{\ell_1-2}(\kappa_2) \\ 0 & 0 & \cdots & 1 & -s_{\ell_1-1} + \pi_{\ell_1-1}(\kappa_1)x & 0 & \cdots & 0 & \pi_{\ell_1-1}(\kappa_2) \\ 0 & 0 & \cdots & 0 & 1-x & -s_{\ell_1} & \cdots & 0 & \pi_{\ell_1}(\kappa_2) \\ \vdots & \vdots & \ddots & \vdots & \vdots & \vdots & \ddots & \vdots & \vdots \\ 0 & 0 & \cdots & 0 & 0 & 0 & \cdots & -s_{\ell_2-2} & \pi_{\ell_2-2}(\kappa_2) \\ 0 & 0 & \cdots & 0 & 0 & 0 & \cdots & 1 & -s_{\ell_2-1} + \pi_{\ell_2-1}(\kappa_2) \end{pmatrix} = 0,$$

where $\pi_i(\kappa_1)$ ($\pi_i(\kappa_2)$) is the numbers of fragments of size $i$ emerged in fragmentation according to the pattern $\kappa_1$ ($\kappa_2$); $\ell_1$ ($\ell_2$) is the size at which the fragmentation occurs in $\kappa_1$ ($\kappa_2$). We choose the order of pure life cycles in a way that $\ell_1 \leq \ell_2$. This determinant contains non-zero elements only at the main diagonal, the first lower subdiagonal; and in $(\ell_1 - 1)$-th and $(\ell_2 - 1)$-th columns. In this case, the characteristic equation is

$$\begin{aligned} p(\lambda, x) &= x p_1(\lambda) R(\lambda) + (1-x) p_2(\lambda), \text{ where} \\ p_1(\lambda) &= F_{\ell_1}(\lambda) - \sum_{i=1}^{\ell_1-1} \pi_i(\mathbf{q}_1) F_i(\lambda), \\ p_2(\lambda) &= F_{\ell_2}(\lambda) - \sum_{i=1}^{\ell_2-1} \pi_i(\mathbf{q}_2) F_i(\lambda), \\ R(\lambda) &= \frac{F_{\ell_2}(\lambda) - \sum_{i=\ell_1}^{\ell_2-1} \pi_i(\mathbf{q}_2) F_i(\lambda)}{F_{\ell_1}(\lambda)}. \end{aligned} \tag{35}$$

The characteristic polynomials $p_1(\lambda)$ and $p_1(\lambda)$ are getting from the pure life cycles $\mathbf{q}_1$ and $\mathbf{q}_2$. Note that if $\ell_1 = \ell_2$ (fragmentation occurs at the same size for both patterns), $R(\lambda) = 1$.

### I Growth rate of a coexisting life cycles in long seasons regime

We consider a pair of fragmentation patterns to be coexisting, if the maximal size of groups allowed by one pattern is smaller than the minimal size of groups produced by another. We call the combination of these two fragmentation patterns a "coexisting life cycle". Then, groups emerging in the larger fragmentation mode, cannot be involved in the fragmentation of the smaller mode. For a group size limit of four cells, there is only one coexisting pair: 1+1 and 2+2. According to Eq. (34), such a mixed life cycle is given by the set of probabilities $\mathbf{q}(x) = (x; 0, 0; 0, 1, 0, 0)$. Groups of the minimal size (one cell) fragment according to the pattern 1+1 with probability $x$, and groups of size larger than 1 fragment according to the pattern 2+2 with probability 1.

To calculate the growth rate of the coexisting life cycle, we start from Eq. (35). In the case of an arbitrary coexisting life cycle, the "larger" fragmentation mode $\mathbf{q}_2$ does not produce groups smaller than $\ell_1$, i.e. $\pi_i(\mathbf{q}_2) = 0$ for all $i < \ell_1$, then the characteristic polynomial $R(\lambda)$ from Appendix H becomes

$$R(\lambda) = \frac{p_2(\lambda)}{F_{\ell_1}(\lambda)}. \tag{36}$$

Therefore, we get

$$p(\lambda, x) = p_2(\lambda)\left(1 - x + \frac{xp_1(\lambda)}{F_{\ell_1}(\lambda)}\right) = \frac{p_2(\lambda)}{F_{\ell_1}(\lambda)}\left(p_1(\lambda) + (1-x)\sum_{i=1}^{\ell_1 - 1}\pi_i(\mathbf{q}_1)F_i(\lambda)\right). \quad (37)$$

The root of the polynomial $p_2(\lambda)$ is always a solution of $p(\lambda, x)$, independently on $x$. In other words, the population always has an option to adopt the pure "larger" life cycle $\mathbf{q}_2$ for any given $x$. The largest root of the second term in Eq. (37), denoted as $\tilde{\lambda}(x)$, is a increasing function of $x$ and is not larger than the largest root of the polynomial $p_1(\lambda)$ (because the additional term is non-negative: $\pi_i(\mathbf{q}_1) \geq 0$, $(1-x) \geq 0$, and $F_i(\lambda) > 0$ for $\lambda > 0$). At $x = 1$, the second term becomes exactly $p_1(\lambda)$, and consequently, $\tilde{\lambda}(1) = \lambda(\mathbf{q}_1)$. The actual growth rate of the population in a coexisting life cycle is the maximum of $\lambda(\mathbf{q}_2)$ and $\tilde{\lambda}(x) \leq \lambda(\mathbf{q}_1)$.

If the season favours pure life cycle $\mathbf{q}_2$ over pure $\mathbf{q}_1$, then $\lambda(\mathbf{q}_1) < \lambda(\mathbf{q}_2)$. Therefore, in the mixed life cycle $\mathbf{q}(x)$, the growth rate will be $\lambda(x) = \lambda(\mathbf{q}_2)$ for any value of $x$. In terms of population behaviour, the population will execute the pure "larger" life cycle, so the value of $x$ is irrelevant. If the season favours pure life cycle $\mathbf{q}_1$ over pure $\mathbf{q}_2$, then $\lambda(\mathbf{q}_1) > \lambda(\mathbf{q}_2)$. Then, the population growth will change depending on $x$. We define $x_0$ as the point, where $\tilde{\lambda}(x_0) = \lambda(\mathbf{q}_2)$. For $x < x_0$, $\lambda(\mathbf{q}_2) > \tilde{\lambda}(x)$, and thus the population growth becomes $\lambda(\mathbf{q}_2)$, similar to the previous case. For $x > x_0$, the population growth becomes $\tilde{\lambda}(x) > \lambda(\mathbf{q}_2)$. It follows that during the season favouring $\mathbf{q}_1$, the largest growth rate is achieved by the life cycle maximizing $\tilde{\lambda}(x)$, which happens at $x = 1$.

Next, we proceed to the dynamic environment and consider a long season regime, where the overall growth rate $\Lambda$ is a weighted sum of growth rates in each of the seasons. If $\mathcal{S}_1$ promotes 1+1 and $\mathcal{S}_2$ promotes 2+2, then the growth in the first season (where $\lambda(\mathbf{q}_{1+1}) > \lambda(\mathbf{q}_{2+2})$) is achieved at $x = 1$ and the growth rate in the second season (where $\lambda(\mathbf{q}_{1+1}) < \lambda(\mathbf{q}_{2+2})$) is independent on $x$. Overall, the optimal life cycle in any such environment is $\mathbf{q}(x)|_{x=1} = (1; 0, 0; 0, 1, 0, 0)$.

### J Long season approximation in near neutral environments for small life cycles $n = 2$

Here, we consider the evolutionarily optimal life cycles of groups not exceeding size two in near neutral environments. Such a population has an access to only three fragmentation patterns: 1+1, 2+1, and 1+1+1. An arbitrary mixed life cycle $\mathbf{q}$ can be characterized by the probabilities of fragmentation according to 1+1 ($u$) and 2+1 ($v$): $\mathbf{q} = (q_{1+1}; q_{2+1}, q_{1+1+1}) = (u; v, 1-v)$. The corresponding projection matrix is (see Eq. (18))

$$A(\mathbf{q}, \mathcal{S}) = \begin{pmatrix} -1 + 2u & 2b_2(v + 3(1-v)) \\ 1 - u & -2b_2(1-v) \end{pmatrix}, \quad (38)$$

with $b_1 = 1$.

In near neutral environment $\mathcal{D} = \{\mathcal{S}_1, \mathcal{S}_2, \tau_1, \tau_2\}$ given by seasons $\mathcal{S}_1 = (1, 1 + \epsilon\beta)$ and $\mathcal{S}_2 = (1, 1 + \epsilon\gamma)$ with season lengths $\tau_1 = Tt/(1+t)$ and $\tau_2 = T/(1+t)$, respectively. The growth rate of a population in the long seasons regime is given by

$$\begin{aligned} \Lambda &\approx \frac{t}{1+t}\lambda(\mathbf{q}, \mathcal{S}_1) + \frac{1}{1+t}\lambda(\mathbf{q}, \mathcal{S}_2) \\ &= 1 + \epsilon\left(\frac{t}{1+t}\lambda'(\mathbf{q}, \beta) + \frac{1}{1+t}\lambda'(\mathbf{q}, \gamma)\right) \\ &+ \frac{\epsilon^2}{2}\left(\frac{t}{1+t}\lambda''(\mathbf{q}, \beta) + \frac{1}{1+t}\lambda''(\mathbf{q}, \gamma)\right) \\ &+ \frac{\epsilon^3}{6}\left(\frac{t}{1+t}\lambda'''(\mathbf{q}, \beta) + \frac{1}{1+t}\lambda'''(\mathbf{q}, \gamma)\right) + O(\epsilon^4). \end{aligned} \quad (39)$$

Note that $\beta$ and $\gamma$ are scalars in this case. A calculation of $\lambda'$, $\lambda''$ and $\lambda'''$ at $\mathcal{S}_1$ gives

$$\begin{aligned}\lambda'(\mathbf{q},\beta) =&\frac{2(1-u)}{5-2u-2v}\beta,\\ \lambda''(\mathbf{q},\beta) =&-\frac{8(1-u)(2-u-v)(3-2v)}{(5-2u-2v)^3}\beta^2,\\ \lambda'''(\mathbf{q},\beta) =&-\frac{48(1-u)(2-u-v)(3-2v)(7+2u(v-2)+v(2v-7))}{(5-2u-2v)^5}\beta^3.\end{aligned} \tag{40}$$

For $\mathcal{S}_2$, the results have the same form with $\gamma$ instead of $\beta$.

If both $\beta$ and $\gamma$ are positive (both negative), the life cycle 2+1 (1+1) is the only evolutionarily optimum in a dynamic environment. Thus, it is worth considering the case of $\beta$ and $\gamma$ having different signs. We assume $\beta > 0$, $\gamma < 0$, because the opposite case will give symmetric results.

First, we consider the local optimality of pure life cycles. At $t \gg 1$, the only local maximum of $\Lambda$ is the pure life cycle 2+1, while at $t \ll 1$, the pure life cycle 1+1 is evolutionary optimal. Supplying Eqs. (40) into Eq. (8), we get

- The life cycle 1+1 is locally stable against small admixture of 2+1 at $t < t_s(1+1,2+1) = -\frac{\gamma}{\beta}$,
- The life cycle 1+1 is locally stable against small admixture of 1+1+1 at $t < t_s(1+1,1+1+1) = -\frac{\gamma}{\beta} - \frac{2}{3}\frac{\gamma}{\beta}(\beta-\gamma)\epsilon + O(\epsilon^2)$,
- The life cycle 2+1 is locally stable against small admixture of 1+1 at $t > t_s(2+1,1+1) = -\frac{\gamma}{\beta} - \frac{8}{81}\frac{\gamma}{\beta}(\beta^2-\gamma^2)\epsilon^2 + O(\epsilon^3)$,
- The life cycle 2+1 is locally stable against small admixture of 1+1+1 at $t > t_s(2+1,1+1+1) = -\frac{\gamma}{\beta} + \frac{1}{3}\frac{\gamma}{\beta}(\beta-\gamma)\epsilon + O(\epsilon^2)$,
- The life cycle 1+1+1 is locally stable against small admixture of 1+1 at $t > t_s(1+1+1,1+1) = -\frac{\gamma}{\beta} - \frac{6}{25}\frac{\gamma}{\beta}(\beta-\gamma)\epsilon + O(\epsilon^2)$,
- The life cycle 1+1+1 is locally stable against small admixture of 2+1 at $t < t_s(1+1+1,2+1) = -\frac{\gamma}{\beta} - \frac{1}{25}\frac{\gamma}{\beta}(\beta-\gamma)\epsilon + O(\epsilon^2)$.

Combining this, the pure life cycle 1+1+1 is never associated to a local maximum of $\Lambda$. The pure life cycle 1+1 is locally stable at $t < t_s(1+1,2+1)$. The pure life cycle 2+1 is locally stable at $t > t_s(2+1,1+1)$. If $|\beta| > |\gamma|$, both the pure life cycle 1+1 and the pure life cycle 2+1 are local maxima of $\Lambda$ at $t$ in the interval between $t_s(2+1,1+1)$ and $t_s(1+1,2+1)$. If $|\beta| < |\gamma|$, there are no evolutionary optimal pure life cycles in this interval of $t$.

Next, we consider the local optimality of mixed life cycles composed of two fragmentation patterns. There are three such combinations:

- The mixed life cycle $\mathbf{q} = (u,1,0)$ composed of 1+1 and 2+1 satisfies $\frac{d}{du}\Lambda = 0$, and therefore is a local extreme, at $t_l = -\frac{\gamma}{\beta} - \frac{\gamma}{\beta}(\beta-\gamma)\frac{4(1-u)u}{(3-2u)^2}\epsilon + O(\epsilon^2)$. This extreme is a local maximum if $\frac{d^2}{du^2}\Lambda < 0$ at this point. The second derivative is $\frac{d^2}{du^2}\Lambda = -\beta\gamma\frac{8(3-4u)}{(3-2u)^5}\epsilon^2 + O(\epsilon^3)$, which is negative at $u > 3/4$. The local maximum is robust against small admixtures of the remaining life cycle 1+1+1 if $\frac{d}{dv}\Lambda > 0$ at this point. The last condition leads to $\frac{d}{dv}\Lambda = -\beta\gamma\frac{4(1-u)}{(3-2u)^3}\epsilon^2 + O(\epsilon^3) > 0$, which is always satisfied for $u \in (0,1)$. Altogether, such a mixed life cycle is evolutionary optimal at $t$ between $t_l|_{u=3/4} = -\frac{\gamma}{\beta} - \frac{1}{3}\frac{\gamma}{\beta}(\beta-\gamma)\epsilon + O(\epsilon^2)$ and $t_l|_{u=1} = -\frac{\gamma}{\beta} + O(\epsilon^2)$.

- The mixed life cycle $\mathbf{q} = (u, 0, 1)$ composed of 1+1 and 1+1+1 satisfies $\frac{d}{du}\Lambda = 0$, and therefore is a local extreme, at $t = -\frac{\gamma}{\beta} - \frac{\gamma}{\beta}(\beta - \gamma)\frac{2(3+2u-2u^2)}{(5-2u)^2}\epsilon + O(\epsilon^2)$. This extreme is a local maximum if $\frac{d^2}{du^2}\Lambda < 0$ at this point. The second derivative is $\frac{d^2}{du^2}\Lambda = -\beta\gamma\frac{11-8u}{(5-2u)^5}\epsilon^2 + O(\epsilon^3)$, however, it is positive for all $u \in (0, 1)$. Therefore, such a mixed life cycle is never an evolutionary optimum.

- The mixed life cycle $\mathbf{q} = (0, v, 1 - v)$ composed of 2+1 and 1+1+1 satisfies $\frac{d}{dv}\Lambda = 0$, and therefore is a local extreme, at $t = -\frac{\gamma}{\beta} - \frac{\gamma}{\beta}(\beta - \gamma)\frac{1-8v+4v^2}{(5-2v)^2}\epsilon + O(\epsilon^2)$. This extreme is a local maximum if $\frac{d^2}{dv^2}\Lambda < 0$ at this point. The second derivative is $\frac{d^2}{dv^2}\Lambda = -\beta\gamma\frac{3-2v}{(5-2v)^5}\epsilon^2 + O(\epsilon^3)$, however, it is positive for all $v \in (0, 1)$. Therefore, such a mixed life cycle is never an evolutionary optimum.

Only one pairwise combination of fragmentation patterns gives rise to the evolutionary optimal mixed life cycle.

Finally, we consider mixed life cycles composed of all three fragmentation patterns. Such a life cycle is an evolutionary optimum if simultaneously $\frac{d}{du}\Lambda = 0$ and $\frac{d}{dv}\Lambda = 0$. If we denote the values of $t$, where these equalities are satisfied by $t_u$ and $t_v$ respectively, it can be shown that $t_u - t_v = -\frac{\gamma}{\beta}(\beta - \gamma)\frac{1}{5-2u-2v}\epsilon + O(\epsilon^2) > 0$. Therefore, it is impossible to find the value of $t$, where both conditions are simultaneously satisfied (i.e. $t_u = t_v$). Therefore, there is no evolutionary optimal mixed life cycles with three components.

Altogether, the areas of optimality of all found life cycles is presented in Fig. 5.

### K Growth rate of the population in near neutral environments is independent on the seasons turnover period $T$

Consider an arbitrary pure or mixed fragmentation mode $\mathbf{q}$ and an arbitrary near neutral dynamic environment $\mathcal{D} = \{\mathcal{S}_1, \mathcal{S}_2, \tau_1, \tau_2\}$, such as the projection matrices in each season are $A_1$ and $A_2$, respectively (see Eq. (18)). In each season, the dynamics of the population within each season is governed by equations

$$\begin{aligned}\dot{\mathbf{x}} &= A_1\mathbf{x},\\ \dot{\mathbf{x}} &= A_2\mathbf{x}.\end{aligned} \tag{41}$$

Let the population composition at the initial moment $\tilde{t} = 0$ be described by the vector of abundances $\mathbf{x}(0)$. Then, the solutions of these equations are

$$\begin{aligned}\mathbf{x}(\tilde{t}) &= e^{A_1\tilde{t}}\mathbf{x}(0),\\ \mathbf{x}(\tilde{t}) &= e^{A_2\tilde{t}}\mathbf{x}(0),\end{aligned} \tag{42}$$

where the matrix exponent is defined as

$$e^A = \sum_{k=0}^{\infty}\frac{A^k}{k!}. \tag{43}$$

After a single cycle of seasons, the population composition is equal to

$$\mathbf{x}(\tau_1 + \tau_2) = e^{A_2\tau_2}e^{A_1\tau_1}\mathbf{x}(0). \tag{44}$$

In the stationary regime of the dynamic environment, the population returns to the same composition after a full round of seasonal changes. However, the population size increases by the factor $e^{\Lambda(\tau_1+\tau_2)}$. Thus,

$$e^{A_2\tau_2}e^{A_1\tau_1}\mathbf{x}(0) = e^{\Lambda(\tau_1+\tau_2)}\mathbf{x}(0). \tag{45}$$

In near neutral environments, cell birth rates are close to one, $b_j = 1+\epsilon\beta_j$ with $\epsilon \ll 1$. Since any projection matrix is linear with respect to cell birth rates $b_j$ (see Eq. (18)), we conclude that projection matrices $A_1$ and $A_2$ can be presented in a form

$$\begin{aligned} A_1 &= A_0 + \epsilon B_1, \\ A_2 &= A_0 + \epsilon B_2, \end{aligned} \tag{46}$$

where $A_0$ is the projection matrix given by fragmentation mode $\mathbf{q}$ and completely neutral environment $b_j = 1$. In the neutral environment, the growth rate of any life cycle is equal to one. Therefore, in the near neutral environment, the growth rate is close to one, i.e.

$$\Lambda = 1 + \epsilon\Lambda_1 + O(\epsilon^2). \tag{47}$$

The evolutionarily optimal life cycle is the one maximizing $\Lambda_1$, so we aim to find this value.

To do that, we use an ansatz for $\mathbf{x}(0)$ in Eq. (45),

$$\mathbf{x}(0) = \mathbf{x}_0 + \epsilon\mathbf{x}_1 + O(\epsilon^2), \tag{48}$$

where $\mathbf{x}_0$ is the right eigenvector of $A_0$ associated with the eigenvalue $\lambda = 1$, i.e. $A_0\mathbf{x}_0 = \mathbf{x}_0$. Plugging Eqs. (46), (47), and (48) into the left hand side of Eq. (45) and discarding terms smaller than $O(\epsilon)$, we have

$$\begin{aligned} e^{A_2\tau_2}e^{A_1\tau_1}\mathbf{x}(0) &\approx e^{(A_0+\epsilon B_2)\tau_2}e^{(A_0+\epsilon B_1)\tau_1}(\mathbf{x}_0+\epsilon\mathbf{x}_1) \\ &= \sum_{k=0}^{\infty}\frac{(A_0+\epsilon B_2)^k\tau_2^k}{k!}\sum_{m=0}^{\infty}\frac{(A_0+\epsilon B_1)^m\tau_1^m}{m!}(\mathbf{x}_0+\epsilon\mathbf{x}_1) \\ &\approx \sum_{k=0}^{\infty}\frac{\tau_2^k}{k!}(A_0^k+\epsilon[A_0^{k-1}B_2+A_0^{k-2}B_2A_0+\ldots+A_0B_2A_0^{k-2}+B_2A_0^{k-1}]) \\ &\times \sum_{m=0}^{\infty}\frac{\tau_1^m}{m!}(A_0^m+\epsilon[A_0^{m-1}B_1+A_0^{m-2}B_1A_0+\ldots+A_0B_1A_0^{m-2}+B_1A_0^{m-1}])(\mathbf{x}_0+\epsilon\mathbf{x}_1) \\ &\approx \sum_{k=0}^{\infty}\sum_{m=0}^{\infty}\frac{\tau_2^k}{k!}\frac{\tau_1^m}{m!}(A_0^{m+k}+\epsilon[A_0^k\{A_0^{m-1}B_1+\ldots+B_1A_0^{m-1}\}+\{A_0^{k-1}B_2+\ldots+B_2A_0^{k-1}\}A_0^m])(\mathbf{x}_0+\epsilon\mathbf{x}_1) \\ &\approx e^{A_0\tau_2}e^{A_0\tau_1}\mathbf{x}_0+\epsilon\left(e^{A_0\tau_2}e^{A_0\tau_1}\mathbf{x}_1+\sum_{k=0}^{\infty}\sum_{m=0}^{\infty}\frac{\tau_2^k}{k!}\frac{\tau_1^m}{m!}R_{km}\mathbf{x}_0\right), \end{aligned} \tag{49}$$

where we defined $R_{km} = A_0^k(A_0^{m-1}B_1+\ldots+B_1A_0^{m-1})+(A_0^{k-1}B_2+\ldots+B_2A_0^{k-1})A_0^m$. Also, plugging Eqs. (46), (47), and (48) into the right hand side of Eq. (45) and discarding terms smaller than $O(\epsilon)$, we have

$$\begin{aligned} e^{\Lambda(\tau_1+\tau_2)}\mathbf{x}(0) &\approx e^{(1+\epsilon\Lambda_1)(\tau_1+\tau_2)}(\mathbf{x}_0+\epsilon\mathbf{x}_1) \\ &\approx e^{\tau_1+\tau_2}(1+\epsilon\Lambda_1(\tau_1+\tau_2))(\mathbf{x}_0+\epsilon\mathbf{x}_1) \\ &\approx e^{\tau_1+\tau_2}\mathbf{x}_0+\epsilon e^{\tau_1+\tau_2}(\Lambda_1(\tau_1+\tau_2)\mathbf{x}_0+\mathbf{x}_1). \end{aligned} \tag{50}$$

Combining Eqs. (49) and (50), we have

$$e^{A_0\tau_2}e^{A_0\tau_1}\mathbf{x}_0+\epsilon\left(e^{A_0\tau_2}e^{A_0\tau_1}\mathbf{x}_1+\sum_{k=0}^{\infty}\sum_{m=0}^{\infty}\frac{\tau_2^k}{k!}\frac{\tau_1^m}{m!}R_{km}\mathbf{x}_0\right) = e^{\tau_1+\tau_2}\mathbf{x}_0+\epsilon e^{\tau_1+\tau_2}(\Lambda_1(\tau_1+\tau_2)\mathbf{x}_0+\mathbf{x}_1). \tag{51}$$

Terms not containing $\epsilon$ cancel each other, because $e^{A_0\tau}\mathbf{x}_0 = e^{\tau}\mathbf{x}_0$, $e^{A_0\tau_2}e^{A_0\tau_1}\mathbf{x}_0 = e^{\tau_1+\tau_2}\mathbf{x}_0$. So, Eq. (51) is reduced to

$$e^{A_0\tau_2}e^{A_0\tau_1}\mathbf{x}_1+\sum_{k=0}^{\infty}\sum_{m=0}^{\infty}\frac{\tau_2^k}{k!}\frac{\tau_1^m}{m!}R_{km}\mathbf{x}_0 = e^{\tau_1+\tau_2}(\Lambda_1(\tau_1+\tau_2)\mathbf{x}_0+\mathbf{x}_1). \tag{52}$$

Next, we multiply Eq. (52) from the left by the left eigenvector of $A_0$, i.e. by the vector $\mathbf{w}_0$, satisfying $\mathbf{w}_0 A_0 = \mathbf{w}_0$ (and $\mathbf{w}_0 e^{A_0\tau} = e^{\tau}\mathbf{w}_0$). Terms containing $\mathbf{x}_1$ cancel each other because $\mathbf{w}_0 e^{A_0\tau_2} e^{A_0\tau_1}\mathbf{x}_1 = e^{\tau_1+\tau_2}(\mathbf{w}_0\mathbf{x}_1)$. Then, Eq. (52) becomes

$$\sum_{k=0}^{\infty}\sum_{m=0}^{\infty}\frac{\tau_2^k}{k!}\frac{\tau_1^m}{m!}\mathbf{w}_0 R_{km}\mathbf{x}_0 = e^{\tau_1+\tau_2}\Lambda_1(\tau_1+\tau_2)(\mathbf{w}_0\mathbf{x}_0), \tag{53}$$

and thus we can find $\Lambda_1$ as

$$\Lambda_1 = \frac{\sum_{k=0}^{\infty}\sum_{m=0}^{\infty}\frac{\tau_2^k}{k!}\frac{\tau_1^m}{m!}\mathbf{w}_0 R_{km}\mathbf{x}_0}{e^{\tau_1+\tau_2}(\tau_1+\tau_2)(\mathbf{w}_0\mathbf{x}_0)}. \tag{54}$$

Now consider the expression $\mathbf{w}_0 R_{km}\mathbf{x}_0$. Since $\mathbf{w}_0$ and $\mathbf{x}_0$ are left and right eigenvectors of $A_0$ associated with unit eigenvalue, then $\mathbf{w}_0 A_0^a B A_0^b \mathbf{x}_0 = \mathbf{w}_0 B\mathbf{x}_0$. Therefore,

$$\begin{aligned}\mathbf{w}_0 R_{km}\mathbf{x}_0 &= \mathbf{w}_0[A_0^k(A_0^{m-1}B_1 + \ldots + B_1 A_0^{m-1}) + (A_0^{k-1}B_2 + \ldots + B_2 A_0^{k-1})A_0^m]\mathbf{x}_0 \\ &= m\mathbf{w}_0 B_1\mathbf{x}_0 + k\mathbf{w}_0 B_2\mathbf{x}_0.\end{aligned} \tag{55}$$

Then, the nominator in Eq. (54) is

$$\begin{aligned}\sum_{k=0}^{\infty}\sum_{m=0}^{\infty}\frac{\tau_2^k}{k!}\frac{\tau_1^m}{m!}\mathbf{w}_0 R_{km}\mathbf{x}_0 &= \mathbf{w}_0 B_1\mathbf{x}_0\sum_{k=0}^{\infty}\sum_{m=0}^{\infty}\frac{\tau_2^k}{k!}\frac{m\tau_1^m}{m!} + \mathbf{w}_0 B_2\mathbf{x}_0\sum_{k=0}^{\infty}\sum_{m=0}^{\infty}\frac{k\tau_2^k}{k!}\frac{\tau_1^m}{m!} \\ &= e^{\tau_1+\tau_2}(\tau_1\mathbf{w}_0 B_1\mathbf{x}_0 + \tau_2\mathbf{w}_0 B_2\mathbf{x}_0)\end{aligned} \tag{56}$$

Plugging this result into Eq. (54), we get

$$\Lambda_1 = \frac{\tau_1\mathbf{w}_0 B_1\mathbf{x}_0 + \tau_2\mathbf{w}_0 B_2\mathbf{x}_0}{(\tau_1+\tau_2)(\mathbf{w}_0\mathbf{x}_0)} = \frac{t\mathbf{w}_0 B_1\mathbf{x}_0 + \mathbf{w}_0 B_2\mathbf{x}_0}{(1+t)(\mathbf{w}_0\mathbf{x}_0)}, \tag{57}$$

where we used $\tau_1 = \frac{tT}{1+t}$ and $\tau_2 = \frac{T}{1+t}$. This shows that in near neutral environments, the growth rate of an arbitrary life cycle $\mathbf{q}$ given by $1+\epsilon\Lambda_1$ depends on the seasons proportion $t$ but is independent on the seasons turnover period $T$.

### L Parameters of presented simulations examples

In this manuscript, we presented a number of optimality maps in various dynamics environments with different combinations of seasons. For the clarity of organisation, here we list birth rates of all dynamic environments illustrated in this paper.

- For the graphs in Fig. 2, we used $\tau_1 = \tau_2 = 2.5$, $\mathcal{S}_1 = (1, 3, 0.5)$, and $\mathcal{S}_2 = (1, 0.5, 3)$.
- For the map in Fig. 3B, we used $\mathcal{S}_1 = (1, 2, 2)$ and $\mathcal{S}_2 = (1, 1, 4)$.
- For the map in Fig. 3C, we used $\mathcal{S}_1 = (1, 3.0, 0.5)$ and $\mathcal{S}_2 = (1, 0.2, 0.5)$.
- For the map in Fig. 3D, we used $\mathcal{S}_1 = (1, 1.5, 1.55)$ and $\mathcal{S}_2 = (1, 0.5, 0.55)$.
- For the map in Fig. 6A, we used $\mathcal{S}_1 = (1, 1.3, 0.95)$ and $\mathcal{S}_2 = (1, 0.95, 1.3)$.
- For the map in Fig. 6B, we used $\mathcal{S}_1 = (1, 4, 0.5)$ and $\mathcal{S}_2 = (1, 0.5, 4)$.
- For the map in Fig. 6C, we used $\mathcal{S}_1 = (1, 0.27, 2.73)$ and $\mathcal{S}_2 = (1, 2.73, 0.27)$.
- For the map in Fig. 6D, we used $\mathcal{S}_1 = (1, 1.101, 2.612)$ and $\mathcal{S}_2 = (1, 0.917, 0.182)$.
